# Peracetyl *N*-cyclobutanoyl-D-mannosamine enhances expression of sialyl-Lewis X (sLeX / CD15s) and adhesion of leukocytes

**DOI:** 10.1101/2021.12.22.473788

**Authors:** Anam Tasneem, Shubham Parashar, Simran Aittan, Tanya Jain, Charu Chauhan, Jyoti Rautela, Zahoor Ahmad Bhat, Kaisar Raza, Arumugam Madhumalar, Srinivasa-Gopalan Sampathkumar

## Abstract

Sialyl-Lewis-X (sLeX/CD15s) epitopes regulate cell adhesion *via* interactions with selectins. High levels of sLeX are associated with chronic inflammation. Intense efforts have been devoted for the development of sLeX mimetics for the prevention of atherosclerosis. By contrast, low levels of sLeX are associated with leukocyte adhesion deficiency (LAD) disorders. In this context, we employed metabolic glycan engineering to alter the fine structures of sialoglycans. Treatment of HL-60 (human acute myeloid leukemia) cells with peracetyl *N*-cyclobutanoyl-D-mannosamine (Ac_4_ManNCb) resulted in a four-fold increase in both sLeX levels and adhesion to E-selectin-Fc chimera. Enhanced sLeX levels on CD162/PSGL-1, CD43, and CD44 were observed through immunoprecipitation. Molecular dynamics (MD) simulations on interactions with E-selectin revealed dramatic differentials in the conformational dynamics of sLeX-Cb compared to sLeX, highlighting the significance of remote *N*-acyl side chains of sialic acids in facilitating bio-active conformations. These results provide opportunities for pharmacological interventions in both LAD and chronic inflammation.

**Table of Contents:** 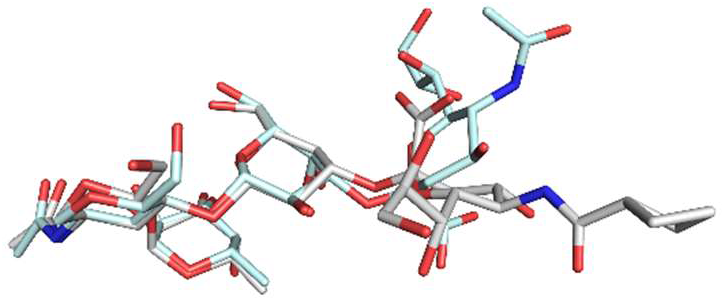

Remote substituents on the tetrasaccharide sialyl-Lewis X (sLeX / CD15s) influence ensembles of bio-active conformations and alter biological outcomes. Treatment with peracetyl *N*-cyclobutanoyl-D-mannosamine (Ac_4_ManNCb) results in four-fold enhancement in the expression of sialyl-Lewis-X (sLeX / CD15s) and adhesion to E-selectin (CD62E). Molecular dynamics simulations show that the sLeX-Cb prefers a flatter bio-active conformation (gray) compared to wild-type sLeX (cyan).

Institute and/or researcher Twitter usernames: @NImmunology @Gops_GlycoIndia

## Introduction

Cell surface glycans, attached to both proteins and lipids, regulate biological processes. Chronic inflammatory diseases, such as atherosclerosis are initiated by dysregulated glycan-endothelial cell interactions. Decades of research have focused on the development of pharmacological mimetics of sialyl-Lewis-X (sLeX; CD15s) for intervention in chronic inflammation^[1]^. On the one hand, enhanced intravascular adhesion of leukocytes and platelets causes inflammatory diseases. On the other hand, defective glycan-protein interactions are responsible for leukocyte adhesion deficiency (LAD) disorders characterized by recurrent infections and poor immune response. It is known that low levels of sLeX in LAD-II results in poor extravasation of neutrophils^[2]^. Enhancing the expression of sLeX through genetic, enzymatic, and pharmacological means in LAD might provide mechanistic insights and therapeutic avenues.

During the past three decades, the ability to modulate glycan structures by exploiting endogenous metabolic pathways has proven to be a powerful tool for study of structure and functions of glycans^[3]^. The application of genetic methods to the study of glycans is complicated by the non-template driven and metabolism dependent biosynthesis. In this context, metabolic glycan engineering (MGE) in living systems provides a much-needed complementary approach to gain insights into glycan-dependent cellular processes in development and diseases^[4]^.

Metabolic processing of *N*-propanoyl-D-mannosamine (ManNProp), a synthetic analogue of the natural substrate *N*-acetyl-D-mannosamine (ManNAc), to *N*-propanoyl-D-neuraminic acid (NeuProp) in rats set the stage for modulation of glycans in living systems^[5]^. Initially, free ManNAc analogues were employed in millimolar concentrations but the efficiency of cellular utilization was improved to micromolar concentrations using the peracetyl derivatives^[6]^. Early studies reported simple hydrocarbon extensions at the *N*-acyl chain of ManNAc and showed their ability to engineer and inhibit glycoconjugate biosynthesis and alter cell-cell, cell-pathogen, and cell-matrix interactions^[7]^. The scope of MGE was expanded with the introduction of chemical reporter groups^[8]^. Bioorthogonal ligations enabled tagging, isolation, imaging, and profiling of glycans through glycomics and glycoproteomics^[3]^. The chemical functionalities presented on ManNAc analogues include ketone, azido, alkynyl, sulfhydryl, diaziridinyl, and cyclopropenyl which undergo orthogonal ligations, respectively, with hydrazide, phosphine / alkyne derivatives, azides, maleimide, light, and tetrazine^[3]^. ManNAc analogues carrying *N*-glycolyl, *N*-phenylacetyl, and halogen substituents at various positions have been reported for their glycan modulating properties^[9]^. Recently, Wittmann and coworkers reported the *N*-cyclopropanoyl-D-mannosamine as a stable variation to the *N*-cyclopropenyl derivative^[10]^.

Although various chemical functional groups have been studied for MGE, the chemical space of *N*-acetyl-D-hexosamine (HexNAc) scaffold is relatively large. The restrictions on chemical structure of ManNAc analogues would only arise due to steric hindrance for processing by the enzymes involved in sialoglycan biosynthesis. In this context, we wanted to explore the *N*-(cycloalkyl)carbonyl derivatives of ManNAc for glycan modulation. We hypothesized that, unlike their straight chain *N*-(alkyl)carbonyl-counterparts, the processing of ManNAc analogues carrying *N*-(cycloalkyl)carbonyl-moieties would be sterically more permissible and result in the expression of corresponding sialic acids with tunable hydrophobicity. The modified sialoglycans might have the potential to alter interactions with sialic acid binding proteins such as selectins, siglecs, and hemagglutinins. The presence of *N*-(cycloalkyl)carbonyl moieties on sialoglycans could potentially modulate the interactions – either *via* direct binding or by altering the bioactive conformations – and hence have the potential to modify cellular properties. Herein, we describe our results on the synthesis of a panel of HexNAc derivatives and their biological evaluation in mammalian cells. Our results show that the ManNAc analogues were readily metabolized and resulted in the enhancement of sialyl-Lewis-X (sLeX / CD15s) epitopes and cell adhesion, in an analogue-structure dependent manner. Molecular dynamics (MD) simulations provided key insights on the importance of remote *N*-acyl moieties on NeuAc of sLeX in altering populations of bio-active conformations.

## Results and Discussion

A panel of peracetylated ManNAc analogues containing *N*-cyclopropanoyl (**1a**, Ac_4_ManNCp)^[10]^, *N*-cyclobutanoyl (**1b**, Ac_4_ManNCb), *N*-azidoacetyl (**1c**, Ac_4_ManNAz)^[11]^, *N*-acetyl (**1d**, Ac_4_ManNAc)^[12]^, *N*-propanoyl (**1e**, Ac_4_ManNProp)^[12]^, *N*-butanoyl (**1f**, Ac_4_ManNBut)^[13]^, and *N*-pentanoyl (**1g**, Ac_4_ManNPent)^[14]^ moieties at the C2-position (**Fig. 1A**) were synthesized. Corresponding *N*-acetyl-D-glucosamine (GlcNAc) analogues, *viz*., *N*-cyclopropanoyl (**2a**, Ac_4_GlcNCp)^[10]^ and *N*-cyclobutanoyl (**2b**, Ac_4_GlcNCb) and peracetyl-*N*-azidoacetyl-D-galactosamine (**3**, Ac_4_GalNAz)^[15]^ were employed as controls.

**Figure 1.**
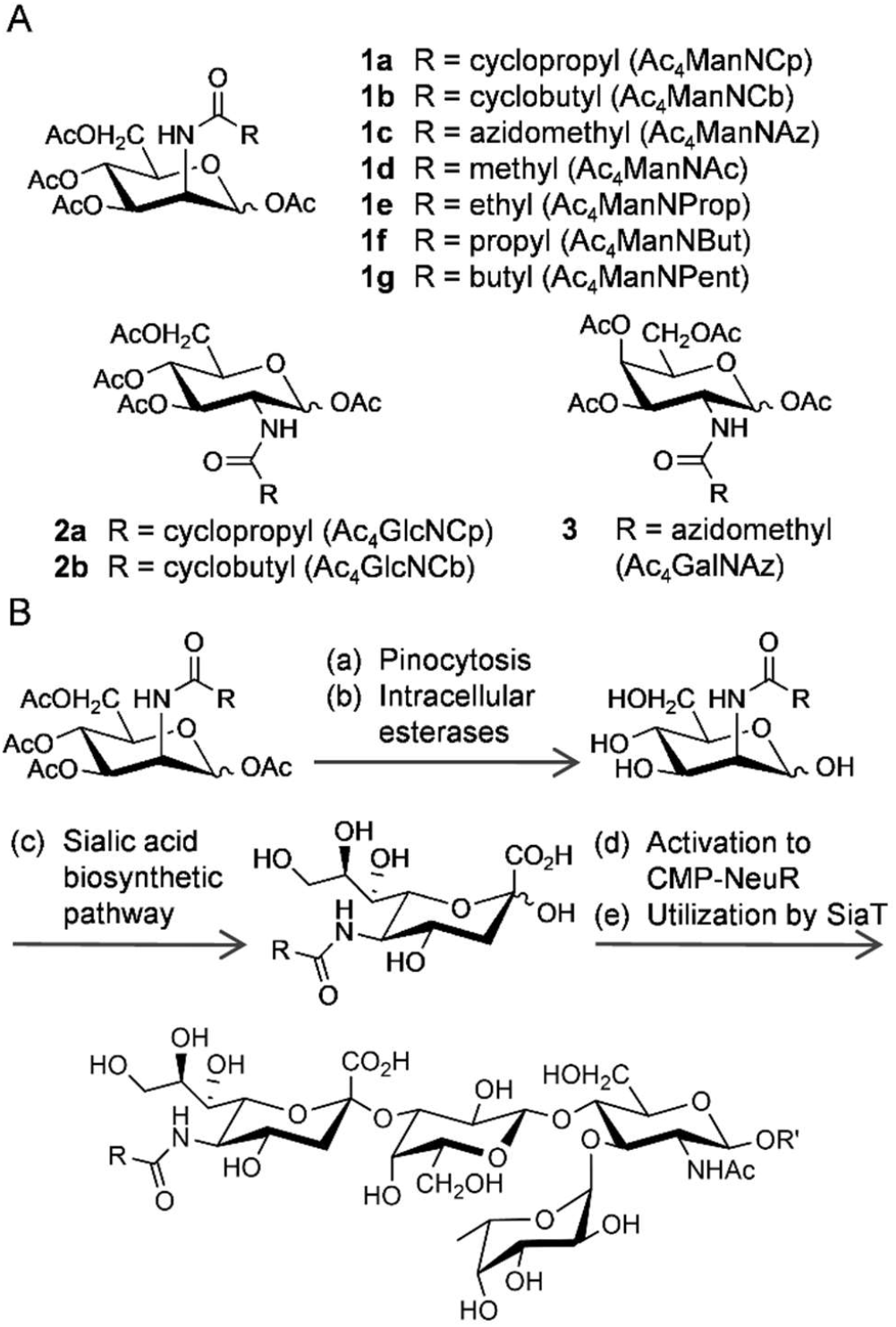
Modulation of sialoglycoconjugates by metabolic processing of *N*-acetyl-D-mannosamine (ManNAc) analogues. (A) Chemical structures of the panel of ManNAc analogues (**1a** – **1g**), GlcNAc analogues (**2a** and **2b**), and a GalNAc analogue (**3**) employed in this study. (B) Metabolic processing of ManNAc analogues through the sialic acid biosynthetic pathway results in the expression of modified sialic acids on *N*-linked and *O*-linked glycoconjugates. Expression of modified sialic acids on sialyl-Lewis-X (sLeX / CD15s) has the potential to alter leukocyte rolling, adhesion, and migration. R’, glycans attached to proteins or lipids; ManNAc, *N*-acetyl-D-mannosamine; GlcNAc, *N*-acetyl-D-glucosamine; GalNAc, *N*-acetyl-D-galactosamine; SiaT, sialyl transferases present in the Golgi; CMP-NeuR, cytidine monophosphate derivative of modified *N*-acetyl-D-neuraminic acid.

Sialic acid biosynthesis begins from the conversion of UDP-GlcNAc to ManNAc by the bifunctional UDP-N-acetyl-D-glucosamine 2-epimerase / *N*-acetyl-D-mannosamine kinase (GNE/MNK) followed reaction with phosphoenol pyruvate to yield *N*-acetyl-D-neuraminic acid (NeuAc). NeuAc is converted to the activated sugar nucleotide donor, CMP-NeuAc, which is then utilized by sialyl transferases stationed in the Golgi apparatus (**Fig. 1B**). Feedback inhibition of GNE/MNK by CMP-NeuAc maintains homeostatic levels of intracellular NeuAc. However, exogenous supply of ManNAc or its analogues could bypass this feedback inhibition and increase the flux of biosynthesis of NeuAc or the corresponding analogues. Depending on the chemical structure, ManNAc analogues are processed to varying degrees of incorporation by the enzymes of the sialic acid pathway.

The ManNAc analogues carrying *N*-(cycloalkyl)carbonyl moieties, **1a** and **1b**, if converted to sialic acids, are not amenable to direct estimation. Hence, we resorted to an indirect approach wherein cells were treated with Ac_4_ManNAz (**1c**) in the presence or absence of **1a** or **1b**. The control (untreated and vehicle treated) cells would express the wild-type NeuAc and the cells treated with **1c** would express both NeuAc and *N*-azidoacetyl-D-neuraminic acid (NeuAz); whereas the cells treated with either **1a** or **1b**, in combination with **1c**, if amenable to metabolism, would result in reduced expression of NeuAz along with expression of either *N*-cyclopropanoyl-D-neuraminic acid (NeuCp) or *N*-cyclobutanoyl-D-neuraminic acid (NeuCb), respectively, in a competitive manner (**Fig. 2A**).

**Figure 2.**
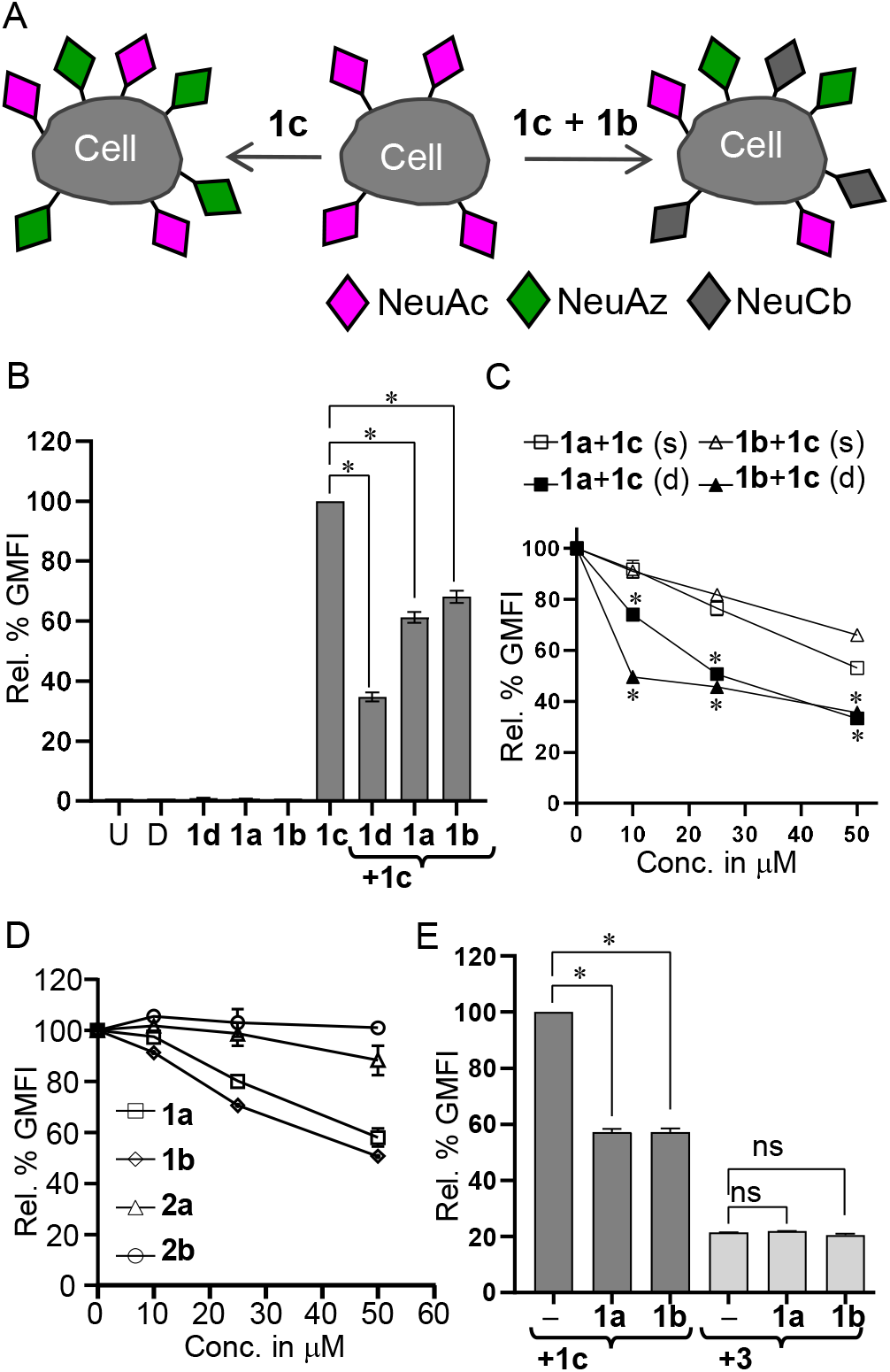
Measurement of processing of ManNAc analogues using Ac4ManNAz as a metabolic reporter in Jurkat cells. (A) Schematic illustration of sialic acids – NeuAc (pink diamonds, wild-type), NeuAz (green), and NeuCb (gray) - expressed on the cell surface. Upon treatment with Ac4ManNAz (**1c**) – either alone or in combination with ManNAc analogues (**1a, 1b, 1d**-**1g**) would alter the expression of NeuAz which could be quantified by SPAAC using DBCO-Cy5 and flow cytometry. (B) NeuAz levels upon incubation with ManNAc analogues – either alone or in combination with **1c** – for 24 h. (C) NeuAz expression in cells treated either simultaneously (**1a** or **1b** with **1c**) or with a delayed addition of **1c** at 12 h, followed by estimation at 36 h. (D) Comparison of GlcNAc analogues (**2a** / **2b**) with ManNAc analogues (**1a** / **1b**) for effect on NeuAz expression by **1c**. (E) Effect of **1a** or **1b** on NeuAz and GalNAz expression, upon co-incubation respectively with **1c** or **3**. Relative GMFI is shown taking NeuAz upon treatment with **1c** alone as 100 % for all the panels. All the analogues were used at a concentration of 50 μM each. Error bars shown are standard deviation of two replicates. Statistical analysis was performed using one-way ANOVA for B and E and two-way ANOVA for C and D, along with Bonferroni’s multiple comparisons test. * *p*-value < 0.05; ns, non-significant. NeuAc, *N*-acetyl-D-neuraminic acid; NeuAz, *N*-azidoacetyl-D-neuraminic acid; NeuCb, *N*-cyclobutanoyl-D-neuraminic acid; U, untreated; D, dimethyl sulfoxide (DMSO, vehicle); Rel. % GMFI, relative percentage geometric mean fluorescence intensity; Conc., concentration, SPAAC, strain promoted azide alkyne cycloaddition; DBCO-Cy5, dibenzocyclooctyne-Cy5.

In order to evaluate the metabolism, we incubated Jurkat (human T-lymphoma) cells with ManNAc analogues either alone (50 μM) or in combination with **1c** (50 μM) for 24 h. Expression of NeuAz on cell surface was estimated by flow cytometry by the strain-promoted azide-alkyne cycloaddition (SPAAC) reaction using dibenzocyclooctyne-Cy5 (DBCO-Cy5)^[16]^. The conditions for click reaction were optimized at various concentrations of DBCO-Cy5 and incubation times (**Supporting Figure 1**). Cells treated with **1c** alone showed robust expression of NeuAz (taken as 100 %), but the NeuAz expression was reduced to 30 %, 60% and 65 % upon co-incubation with **1d, 1a**, and **1b** respectively. Untreated cells and cells treated with vehicle (dimethyl sulfoxide (D)), **1a, 1b**, or **1d** alone showed negligible signals (**Fig. 2B** and **Supporting Figure 2**).

The inhibitory effects on NeuAz expression by **1a** and **1b** were found to be both competitive and dose dependent. Jurkat cells were first treated with either **1a** or **1b** (0 – 50 μM) followed by addition of **1c** (50 μM) at 12 h and estimation of NeuAz at 36 h. The suppression of NeuAz expression was substantially higher when added with a 12 h delay compared to simultaneous addition. At 10 μM, delayed addition resulted in 30% and 50% suppression upon treatment with **1a** and **1b**, respectively; by contrast simultaneous addition resulted only in a 10% reduction of NeuAz for both **1a** and **1b** (**Fig. 2C** and **Supporting Figure 2**). These results suggested that both **1a** and **1b** are amenable to sialic acid biosynthesis and that **1b** showed better competitive ability compared to **1a**. In order to test the specificity of ManNAc analogues, Jurkat cells were incubated with **1a, 1b, 2a**, or **2b** (0 – 50 μM) in combination with **1c** (50 μM, 24 h) (**Fig. 2D**). Unlike **1a** or **1b**, the GlcNAc analogues **2a** or **2b**, the C-2 epimers, were found to cause no inhibition of NeuAz expression, thus affirming the necessity to bypass the GNE/MNK step.

The high degree of selectivity of ManNAc analogues for sialic acid pathway was further confirmed by competitive studies using Ac_4_GalNAz (**3**) which is known to be processed through both the mucin-type *O*-glycosylation (MTOG) and β-*O*-GlcNAc-ylation pathways^[15, 17]^. The expression of NeuAz by **1c** was suppressed by **1a** or **1b** upon co-incubation; by contrast, no reduction in the expression of GalNAz on the cell surface was noticed upon co-incubation of **3** with **1a** or **1b** (**Fig. 2E** and **Supporting Figure 2**). These results confirmed that the ManNAc analogues did not interfere with the GalNAz processing through MTOG.

Having confirmed metabolic acceptability in Jurkat cells, we tested the analogues on HL-60 (human acute myeloid leukemia) cells which are a reliable model for neutrophil adhesion^[18]^. HL-60 cells were treated with **1c** either alone (50 μM) or in combination with ManNAc analogues (50 μM) for 24 h followed by DBCO-Cy5 ligation. Flow cytometry analysis showed that **1a, 1b, 1d, 1e, 1f**, and **1g** reduced the NeuAz expression, respectively, to 62 %, 60 %, 38 %, 70 %, 88 %, and 95 % with respect to **1c** alone (taken as 100 %) (**Fig. 3A**). Maximum suppression was observed for the *N*-acetyl (wild-type) analogue **1d**, followed by the cycloalkyl derivatives **1a** and **1b**. The straight chain analogues **1e, 1f**, and **1g** showed decreasing NeuAz suppression as a function of *N*-alkyl chain length, which was in contrast to the *N*-cycloalkyl moieties. While **1b** induced a reduction of 40 %, the corresponding straight chain analogue with five carbons **1g** showed only a mild reduction of 5.0 % with respect to controls. Similarly, **1a** induced a reduction of 40% compared to only a 15 % reduction by **1f**. These results demonstrated that the cycloalkyl groups at the *N*-acyl chain of ManNAc are significantly more acceptable for metabolic processing compared to their straight chain analogues.

**Figure 3.**
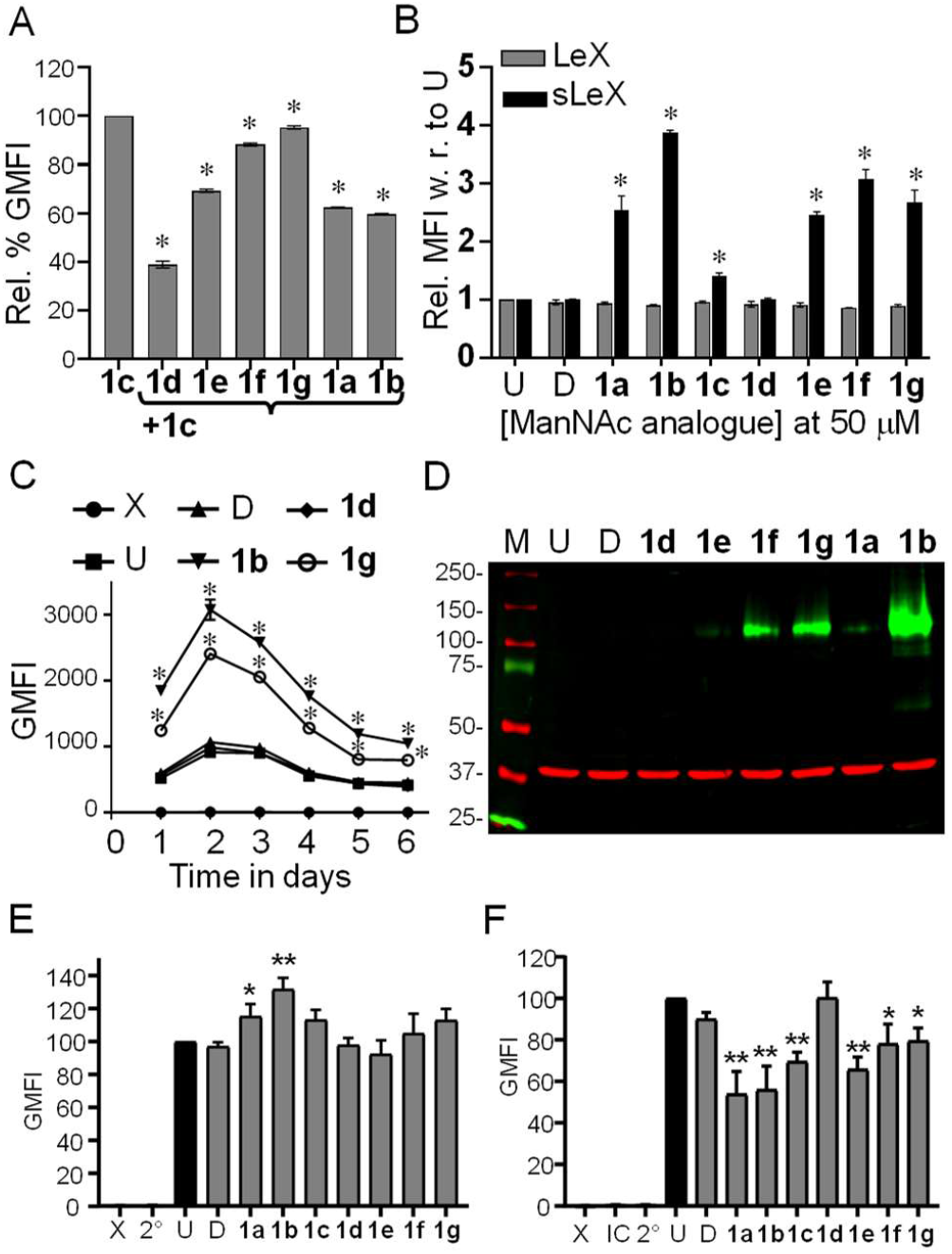
Enhanced expression of sLeX epitopes induced by the *N*-cyclobutanoyl ManNAc analogue 1b in HL-60 cells. (A) Flow cytometry estimation of NeuAz expression upon co-incubation of ManNAc analogues in combination with **1c**. (B) Effect of ManNAc analogues on the cell surface levels sLeX and LeX epitopes measured using anti-CD15s (clone CSLEX1) and anti-CD15 (clone HI98) antibodies, respectively. (C) Time course of the expression of sLeX upon incubation with ManNAc analogues over a period of six days. (D) Cells were incubated with ManNAc analogues for 72 h, lysed, and probed through dual western blots using anti-CD15s (CSLEX1) (upper green bands) and anti-β-actin (lower red bands; loading control). Blots shown are representative of two biological replicates. Flow cytometry estimation of binding of (E) E-selectin-Fc chimera and (F) anti-CLA (HECA452) antibody to cells incubated with ManNAc analogues. All the analogues were used at a concentration of 50 μM each for 24 h, unless stated otherwise. Error bars shown are standard deviation of at least two replicates. Statistical analysis was performed using one-way ANOVA with Bonferroni’s multiple comparisons test for A, E, and F and using paired ‘t’ test for B and C. \**p*-value < 0.001 for A and < 0.05 for B, C, E, and F; ns, non-significant. CLA, cutaneous lymphocyte antigen; M, markers; X, unstained; U, untreated; D, DMSO (vehicle) treated; IC, isotype control; 2°, secondary antibody only; GMFI, geometric mean fluorescence intensity.

Profiling of global changes to glycosylation by far-western blotting and flow cytometry using lectins revealed that treatment with **1b** resulted in a moderate gain in binding to *Maackia amurensis* lectin-II (MAL-II) (which bind to the NeuAcα2→3Gal moiety, a part of sLeX) while a moderate decrease in binding was observed for *Sambucus nigra* agglutinin (SNA) (**Supporting Figure 3**). Notably, cell densities measured at 24, 48, and 72 h, as a measure of proliferation, upon incubation with **1a** – **1g, 2a**, and **2b** (100 μM) exhibited negligible toxicity in all cases (**Supporting Figure 4**).

With the ability to modulate the sialic acids using *N*-(cycloalkyl)carbonyl analogues we ventured to investigate changes to established sialo-epitopes of immunological importance, *viz*., sialyl-Lewis-X (sLeX, CD15s) using anti-CD15s (CSLEX1) antibody. Optimal conditions for flow cytometry were obtained by titration at various concentrations of anti-CD15s. Consistent with the literature, HL-60 cells were positive and Jurkat cells were negative for CSLEX1 epitopes (**Supporting Figure 5**). Strikingly, HL-60 cells upon incubation with **1b** (50 μM, 24 h) showed nearly a four-fold increase in the binding of CSLEX1 antibody, compared to controls (untreated, vehicle-treated, and **1d**) (**Fig 3B**). Treatment with **1f** resulted in a three-fold increase; the three analogues, *viz*., **1a, 1e**, and **1g** resulted in an increase of ∼2.5-fold while **1c** showed a moderate increase of 1.4-fold. Notably, *N*-butanoyl analogue **1f** showed higher CSLEX1 binding compared to the four-carbon *N*-cyclopropanoyl analogue **1a**; whereas the *N*-pentanoyl analogue **1g** showed lower CSLEX1 binding compared to the five-carbon *N*-cyclobutanoyl analogue **1b**. Levels of LeX/CD15 levels, measured using the anti-CD15 (HI98) antibody, remained unaffected upon treatment with **1a** – **1g** (**Fig. 3B**), suggesting that there has been no trade-off in the LeX levels for sLeX. The enhanced binding of anti-sLeX antibody is likely due to the expression of modified sialoglycans rather than a net increase in sLeX (wild-type) levels, since **1d** did now show any effect. These results suggested unique analogue structure-dependent differential effects of the *N*-acyl moiety for metabolic processing and sLeX display.

Time course studies revealed that incubation of HL-60 cells with a single dose of **1b** or **1g** resulted in peak sLeX expression on day 2 and returned to normal levels by day 6. There was a maximum of 3.0-fold and 2.2-fold increase in sLeX levels upon treatment, respectively, with **1b** and **1g** on day 2 while no change was observed in cells treated with controls (vehicle and **1d**). By day 6, cells treated with **1g** showed a slightly faster recovery to native levels while **1b** treatment still maintained approximately 2.3-fold higher levels of sLeX (**Fig. 3C**). These results confirmed the advantages of pharmacological approaches, with dosage control and reversibility, over genetic methods for modulation of sLeX.

Total sLeX levels in HL-60 cells incubated with ManNAc analogues were probed by western blotting. Robust increase in sLeX bands, stained using CSLEX1 antibody, were observed between 150-100 kDa range upon treatment for **1b**, with moderate increase for **1f** and **1g**, and marginal increase for **1a** and **1e** (**Fig. 3D**). Controls (U, D, or **1d**) were negative for sLeX epitopes, consistent with earlier reports on spontaneous differentiation of HL-60 cells to hyposialylated phenotypes^[19]^. Earlier reports have shown enhanced expression of sLeX epitopes (probed with KM93 antibody) in HL-60 cells upon treatment with ManNProp at 10 mM^[20]^. Only a mild increase in sLeX was observed with its peracetylated counterpart **1e** at 50μM. While **1f** showed an increase in sLeX levels in HL-60 cells, the corresponding *N*-cyclopropanoyl analogue **1a** showed only a marginal increase. By contrast, the *N*-cyclobutanoyl analogue **1b** showed maximum increase in sLeX as compared to the five-carbon straight chain analogue **1g** (**Fig. 3D**). These results showed the unique properties of the *N*-cyclobutanoyl derivative **1b**, which was not observed with the *N*-cyclopropanoyl derivative **1a**, thus highlighting the significance of ManNAc analogue-structure dependent effects on cellular outcomes. The increase in sLeX levels upon treatment with **1b** could be attributed to two plausible reasons, *viz*., (i) enhanced biosynthesis of both wild-type and modified NeuCb carrying sLeX (sLeX-Cb) and (ii) enhanced binding affinity of CSLEX1 antibody to sLeX-Cb.

Next, we tested the effects on the binding of sLeX to its endogenous receptor E-selectin (CD62E). Binding of both mouse and human E-selectin-Fc chimera protein to HL-60 cells were studied under optimal conditions (**Supporting Figure 6**). Consistent with earlier reports, mouse E-selectin-Fc (mCD62E) was found to show higher binding to HL-60 cells compared to human E-selectin-Fc (hCD62E)^[21]^. Flow cytometry results revealed a robust increase of 2.3-fold in cells treated with **1b** with respect to controls, similar to the results obtained for CSLEX1 binding (**Fig. 3E**). Cells treated with vehicle (D), **1d**, or **1a** showed no significant change with respect to untreated controls.

Cutaneous lymphocyte antigens (CLA) carry sLeX glycans, recognized by the anti-CLA antibody (HECA452)^[22]^, which interact with E-selectin and enable T-cell migration. We tested the effect of ManNAc analogues treatment on HECA binding in HL-60 cells. Surprisingly, a prominent 40% decrease in HECA-452 binding was observed upon treatment with **1a** or **1b** (50 μM, 24 h) compared to untreated cells; while cells treated with vehicle (D) or **1d** showed only a moderate increase (**Fig. 3F**). While a robust increase was found for binding of mCD62E and CSLEX1, a decrease in HECA452 was observed. A possible explanation is that HECA452 is specific for human sLeX carrying NeuAc and does not bind to mouse sLeX carrying NeuGc, suggesting requirement for direct contacts with the *N*-acyl moiety for binding^[23]^. Similarly, in the case of **1a** or **1b**, it is possible that the HECA452 binding is abrogated, respectively, due to presence of NeuCp or NeuCb on sLeX. Also, increased sLeX biosynthesis, as a consequence of treatment with **1a** or **1b**, could in turn affect both glycan site occupancy and glycan densities. Since HECA-452 is a non-function blocking antibody, it might depend partially on accessibility of the polypeptide backbone for contacts which could be masked by enhanced glycan site occupancy and the presence of sLeX-Cb.

Next, we focused our attention to characterizing sLeX on specific glycoproteins. Many glycoproteins, including CD162/PSGL-1 (P-selectin glycoprotein ligand-1), CD43, CD44, and ESL-1, have been reported to carry sLeX^[24]^. HL-60 cells were treated with ManNAc analogues for 72 h, lysed, and subjected to immunoprecipitation using anti-CD162/PSGL-1 (KPL-1), anti-CD43 (C-20), and anti-C44 (IM7) antibodies and probed for sLeX with CSLEX1 antibody (**Fig. 4A** and **Supporting Figure 7**). Cells treated with vehicle (D), **1c**, or **1d** were negative for sLeX epitopes on CD162/PSGL-1 similar to untreated cells; **1a** and **1e** showed moderate sLeX levels; intense bands were observed for samples treated with **1b, 1f**, and **1g**. CD43 immunoprecipitates showed faint bands for **1a** and **1g**, moderate levels for **1f**, and maximal levels for **1b**; D, **1c, 1d, 1e** were negative similar to untreated cells. CD44 immunoprecipitates showed robust expression of sLeX levels for **1b** and moderate levels for **1g**, with controls (vehicle and **1d**) being negative. These results highlight the non-overlapping structure-dependent effect of ManNAc analogues on CD antigens and that **1b** was able to enhance sLeX epitopes on all the three glycoproteins.

**Figure 4.**
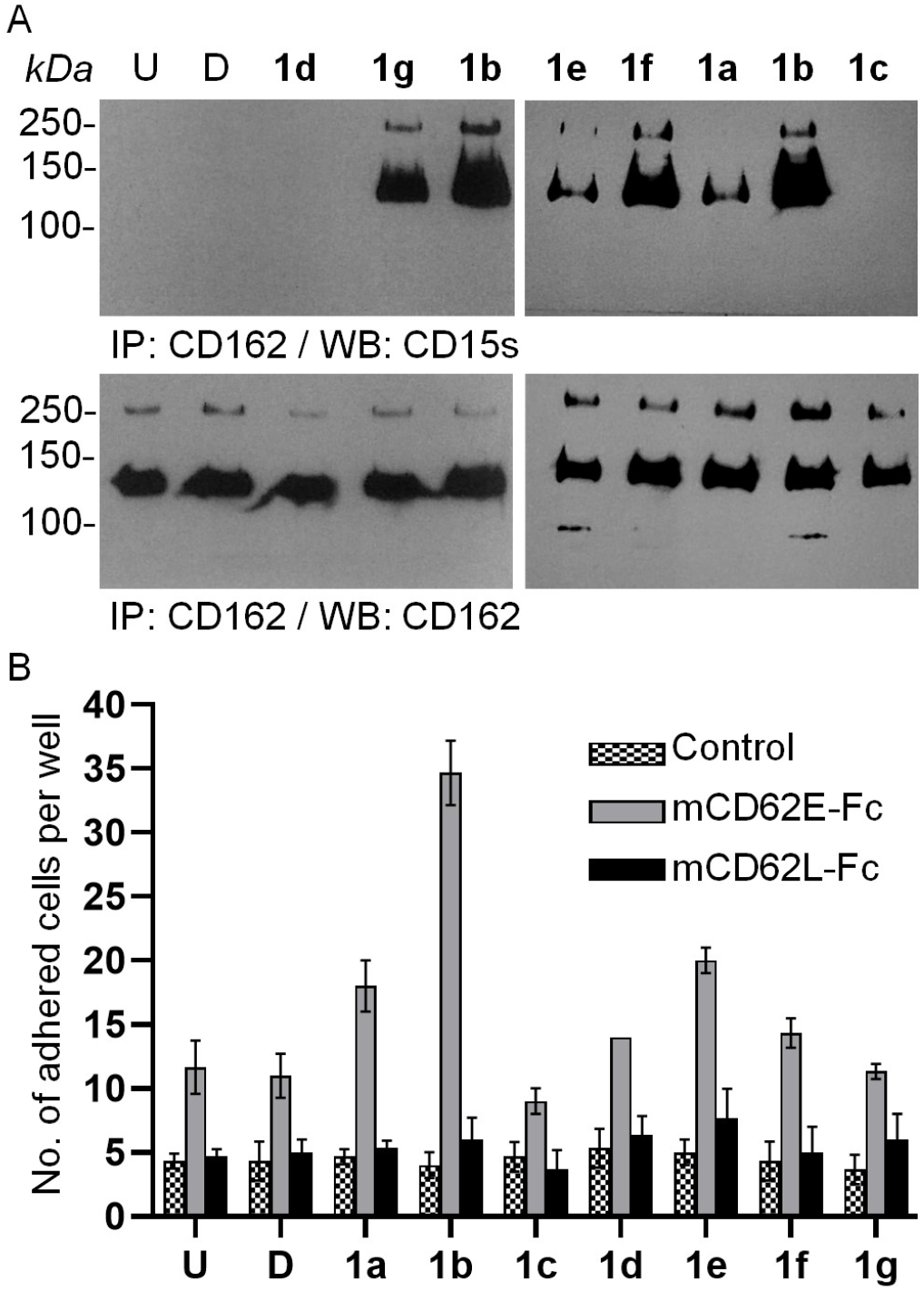
ManNAc analogues increase sLeX on CD162/PSGL-1 and facilitate cell adhesion. A) HL-60 cells were incubated with **1a** – **1g** (50 μM) along with vehicle (D) and untreated controls for 72 h. Cells lysates were subjected to immunoprecipitation using anti-CD162/PSGL-1 (clone KPL-1) and the immunoprecipitates were resolved on SDS-PAGE, blotted onto nitrocellulose membranes, and probed using anti-CD15s (CSLEX1) (upper panel) and anti-CD162/PSGL-1 (lower panel). Blots shown are representative of at least two biological replicates. B) Number of HL-60 cells, pre-incubated with **1a** – **1g** (50 μM, 48 h), adhered to surfaces coated with either mCD62E-Fc chimera or mCD62L-Fc chimera on protein-G and blocked with BSA. Surfaces coated only with protein-G and BSA were employed as controls; adhered cells were fixed, labeled with DAPI, and enumerated using fluorescence microscopy; two field views were obtained for each well in 96-well plates. Error bars shown are standard deviation of at least three replicates for each condition. Statistical analysis was performed using two-way ANOVA with Bonferroni’s multiple comparisons test. * *p*-value < 0.05; ns, non-significant.

Having established the unique properties of **1b** in enhancing the expression of sLeX, we turned our attention to functional studies. Cell adhesion studies were performed on plates coated with protein-G followed by either mouse E-selectin-Fc (mCD62E) or mouse L-selectin-Fc (mCD62L) chimera, followed by blocking with BSA. Plates coated with only protein-G and BSA were used as controls. L-selectin was included in order to study the effect of ManNAc analogues on sulfo-sLeX expression^[25]^. HL-60 cells were first incubated with the ManNAc analogues **1a** – **1g** for 48 h and then allowed to adhere to selectin-coated surfaces. The adhered cells were fixed, stained with DAPI, imaged by fluorescence microscopy, and the numbers of cells were enumerated (**Fig. 4B** and **Supporting Figure 8**). Treatment with **1b** resulted in a robust 3.5-fold increase in the numbers of cells adhered on mCD62E compared to controls (untreated and vehicle treated); **1a** and **1e** showed a 1.5-fold increase while **1c, 1d, 1f**, and **1g** showed negligible effects. Strikingly, while treatment with **1b** enhanced cell attachment the corresponding straight chain analogue **1g** did not show any effect. No differences were observed for mCD62L and controls under all conditions indicating the low abundance of sulfo-sLeX on HL-60 cells. Clearly, **1b** displayed unique ability to enhance sLeX levels on glycoproteins, including CD162, CD43, and CD44, as shown by CSLEX1 and MAL-II and resulted in increased number of cells attached to E-selectin coated surfaces.

In order to gain mechanistic insight, we resorted to MD simulation studies of sLeX-E-selectin interactions. Earlier MD simulations focused on design of minimal ligands and on the conformational flexibility of sLeX^[26]^. Theoretically, the enhanced binding, induced by **1b**, could be the result of direct contacts of the *N*-cyclobutanoyl moiety with hydrophobic residues on CSLEX1 and E-selectin. Alternatively, the remote *N*-acyl substituents on NeuAc might influence and favor populations of bio-active conformations of sLeX. The paratopes of CSLEX1 are not known. However, several crystal structures of E-selectin-sLeX complexes are available^[27]^. Detailed structural studies have established that E-selectin binding to sLeX is determined by the 3-OH/4-OH of the fucosyl residue and the carboxylate moiety of NeuAc with the *N*-acetyl group projecting out to the solvent ^[1d]^.

E-selectin consists of lectin domain, epidermal growth factor-like (EGF-like) domain, and short consensus repeats 1 and 2 (SCR1 and SCR2) connected by a linker, that exhibit spring-like action^[27]^. We choose the coordinates of E-selectin (PDB: 1G1T) for MD simulation in complex with either sLeX or sLeX-Cb (**Supporting movies 1-4**). Simulations of sLeX and sLeX-Cb as unbound ligands showed flexible conformations occupying various dihedral angles for the tetrasaccharides (**Supporting Figure 9-14 and Supporting Tables 1-4**). Striking differences were noted for the preferred conformations of E-selectin bound ligands highlighted by the dihedral angles (**Fig. 5A**). NeuAcα2→3Gal preferred −44 and −143 for Φ and Ψ dihedral values, respectively, while the NeuCbα2→3Gal preferred +79 and −115. Major populations of sLeX displayed a S-shaped (bent) conformation while sLeX-Cb displayed a J-shaped (straight) conformation (**Fig. 5B** and **5C**). No major changes were observed for Fucα1→3GlcNAcβ1→4Gal moiety. Dwell times showed that the interaction of glycerol side chain of NeuAc with E-selectin was dominant for sLeX (**Fig. 5D**). By contrast the interaction of carboxylate chain of NeuAc with E-selectin was dominant for sLeX-Cb (**Fig. 5E**). Consistent with the higher binding energy of Coulomb interactions and the criticality of the carboxylate moiety, the sLeX-Cb displayed higher proportion of bioactive conformational populations compared to sLeX. Interestingly, sLeX binding necessitated energy expending stretching movement of the lectin and EGF-like domains through the linker^[28]^; whereas, sLeX-Cb preferred binding to the unstretched (resting) state of E-selectin (**Supporting information**). The insights provided by MD simulation on the differential binding properties induced by remote substituents on ligands and receptor dynamics might have far-reaching implications for carbohydrate-based drug design.

**Figure 5.**
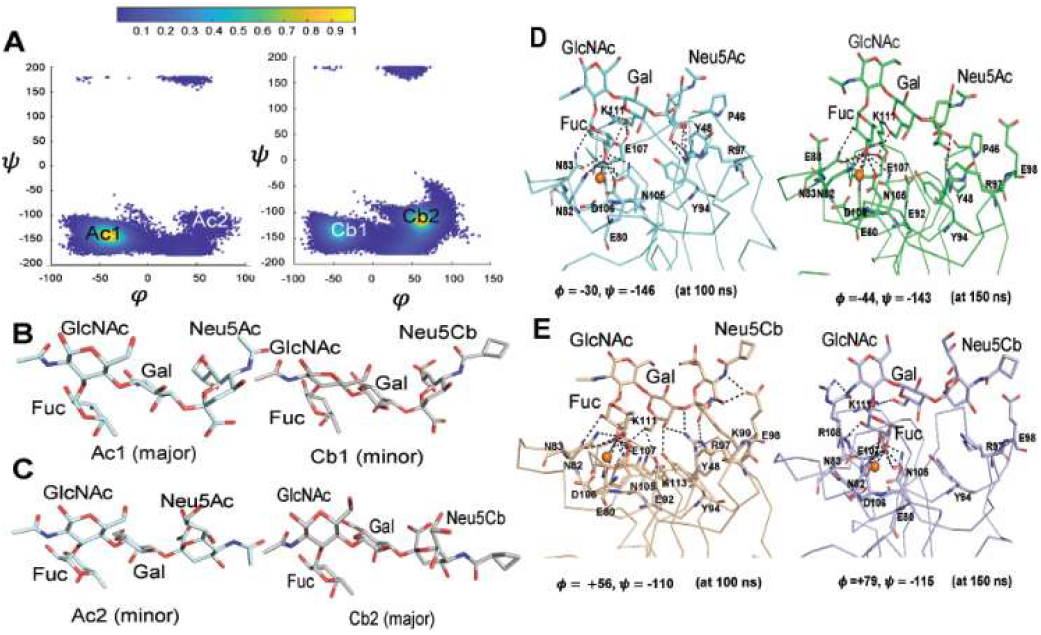
Molecular dynamics (MD) simulations revealed altered populations of bioactive conformations. A) Dihedral plots for NeuAcα2→3Gal conformations of sLeX (200 ns) and sLeX-Cb (300 ns), bound to E-selectin (PDB: 1G1T); coordinates for sLeX were obtained from the GlyCAM website (*see* **Supporting Information** for movies on MD simulation). Major and minor conformers for (B) sLeX and (C) sLeX-Cb are shown. Representative view of bound conformers, at 100 and 150 ns, shown for (D) sLeX and (E) sLeX-Cb ; the dihedral angles for NeuAcα2→3Gal glycosidic bond are shown.

## Conclusion

Sialoglycans are critically involved in the regulation of cellular immune homeostasis through the sLeX-selectin (E/L/P) axis^[29]^ for cell-cell interactions, cell adhesion, and cell migration^[30]^. On the one hand, decades of research efforts have been devoted to the development of pharmacological agents that could inhibit sLeX-selectin interactions under inflammatory conditions^[1d]^; On the other hand, disease conditions such as LAD^[2b]^ are highlighted by the low levels of sLeX and poor leukocyte adhesion to selectins. In this context our investigations with a panel of HexNAc analogues revealed that the *N*-cyclobutanoyl ManNAc analogue, **1b**, readily enhanced sLeX levels and resulted in enhanced adhesion to E-selectin. MD simulations studies provided insights on the influence of remote *N*-acyl substituents in favoring bioactive conformations. We have shown the analogue-structure dependent effects on sLeX expression and consequential modulation of their binding to anti-glycan antibodies and lectins. The ability to either enhance or suppress sLeX-selectin^[18, 31]^, sialoglycans-siglec^[29a]^, and sialoglycans-viral hemagglutinin interactions^[32]^ using pharmacological agents has the potential to open up new avenues for therapeutics for immune deficiency disorders as well as shed light on the intricate mechanisms of functional roles of glycans in immune regulation.

## Experimental Section

Supporting results and discussion, supporting figures, materials and methods for cell culture experiments, and chemical synthesis, NMR spectra, and methods along with movies for MD simulation are provided in the Supporting Information.

## Supporting information

Supporting Movie 1_sLeX-Cb

Supporting Movie 2_sLeX

## Acknowledgements

Funding support from Government of India, Department of Science and Technology (DST) NanoMission grant No. SR/NM/NB-1093/2017(G) to both SGS and KR, DST Science & Engineering Research Board (DST-SERB) grant No. EMR/2016/008000 to SGS, and NII intramural grants to SGS are gratefully acknowledged. AT and SP were supported by fellowships from Department of Biotechnology (DBT), DBT/2017/NII/845, and Indian Council of Medical Research (ICMR), F. No. 45/52/2018-NAN/BMS, respectively. We thank Dr. Devinder Sehgal for access to the fluorescent microscope. We thank Ms. Shanta Sen (Mass Spectrometry), Mr. T. Khaling (Flow Cytometry), and Mr. Kevla Nand and Mr. Shahnawaz Haider (Instrumentation Facility) at NII. Technical help from Mr. Mohd. Aslam, Mr. Anand Kumar Toppo, and Mr. P. Rajkumar are gratefully acknowledged.

## References

[1] a) F. Jin, F. Wang, Glycoconj J 2020, 37, 277–291; b) M. L. Phillips, E. Nudelman, F. C. Gaeta, M. Perez, A. K. Singhal, S. Hakomori, J. C. Paulson, Science 1990, 250, 1130–1132; c) G. Walz, A. Aruffo, W. Kolanus, M. Bevilacqua, B. Seed, Science 1990, 250, 1132–1135; d) J. L. Magnani, Arch Biochem Biophys 2004, 426, 122–131.

[2] a) D. J. Becker, J. B. Lowe, Biochim Biophys Acta 1999, 1455, 193–204; b) Y. Katayama, A. Hidalgo, J. Chang, A. Peired, P. S. Frenette, J Exp Med 2005, 201, 1183–1189.

[3] C. Agatemor, M. J. Buettner, R. Ariss, K. Muthiah, C. T. Saeui, K. J. Yarema, Nat Rev Chem 2019, 3, 605–620.

[4] M. J. Buettner, S. R. Shah, C. T. Saeui, R. Ariss, K. J. Yarema, Front Immunol 2018, 9, 2485.

[5] H. Kayser, R. Zeitler, C. Kannicht, D. Grunow, R. Nuck, W. Reutter, J Biol Chem 1992, 267, 16934–16938.

[6] E. J. Kim, S. G. Sampathkumar, M. B. Jones, J. K. Rhee, G. Baskaran, S. Goon, K. J. Yarema, J Biol Chem 2004, 279, 18342–18352.

[7] a) O. T. Keppler, R. Horstkorte, M. Pawlita, C. Schmidt, W. Reutter, Glycobiology 2001, 11, 11R–18R; b) S. Goon, B. Schilling, M. V. Tullius, B. W. Gibson, C. R. Bertozzi, Proc Natl Acad Sci U S A 2003, 100, 3089–3094.

[8] J. A. Prescher, C. R. Bertozzi, Cell 2006, 126, 851–854.

[9] C. D. Rillahan, A. Antonopoulos, C. T. Lefort, R. Sonon, P. Azadi, K. Ley, A. Dell, S. M. Haslam, J. C. Paulson, Nat Chem Biol 2012, 8, 661–668.

[10] J. Hassenruck, V. Wittmann, Beilstein J Org Chem 2019, 15, 584–601.

[11] E. Saxon, C. R. Bertozzi, Science 2000, 287, 2007–2010.

[12] C. L. Jacobs, S. Goon, K. J. Yarema, S. Hinderlich, H. C. Hang, D. H. Chai, C. R. Bertozzi, Biochemistry 2001, 40, 12864–12874.

[13] L. K. Mahal, N. W. Charter, K. Angata, M. Fukuda, D. E. Koshland, Jr., C. R. Bertozzi, Science 2001, 294, 380–381.

[14] O. T. Keppler, P. Stehling, M. Herrmann, H. Kayser, D. Grunow, W. Reutter, M. Pawlita, J Biol Chem 1995, 270, 1308–1314.

[15] H. C. Hang, C. Yu, D. L. Kato, C. R. Bertozzi, Proc Natl Acad Sci U S A 2003, 100, 14846–14851.

[16] a) N. J. Agard, J. A. Prescher, C. R. Bertozzi, J Am Chem Soc 2004, 126, 15046–15047; b) E. M. Sletten, C. R. Bertozzi, Angew Chem Int Ed Engl 2009, 48, 6974–6998.

[17] a) D. H. Dube, J. A. Prescher, C. N. Quang, C. R. Bertozzi, Proc Natl Acad Sci U S A 2006, 103, 4819–4824; b) M. Boyce, I. S. Carrico, A. S. Ganguli, S. H. Yu, M. J. Hangauer, S. C. Hubbard, J. J. Kohler, C. R. Bertozzi, Proc Natl Acad Sci U S A 2011, 108, 3141–3146.

[18] S. S. Wang, V. D. Solar, X. Yu, A. Antonopoulos, A. E. Friedman, K. Agarwal, M. Garg, S. M. Ahmed, A. Addhya, M. Nasirikenari, J. T. Lau, A. Dell, S. M. Haslam, S. G. Sampathkumar, S. Neelamegham, Cell Chem Biol 2021.

[19] O. T. Keppler, S. Hinderlich, J. Langner, R. Schwartz-Albiez, W. Reutter, M. Pawlita, Science 1999, 284, 1372–1376.

[20] R. Horstkorte, K. Rau, W. Reutter, S. Nohring, L. Lucka, Exp Cell Res 2004, 295, 549–554.

[21] Z. Ni, B. Walcheck, Immunol Lett 2007, 108, 179–182.

[22] C. M. Kummitha, V. S. Shirure, L. F. Delgadillo, S. P. Deosarkar, D. F. Tees, M. M. Burdick, D. J. Goetz, J Immunol Methods 2012, 384, 43–50.

[23] J. Mitoma, T. Miyazaki, M. Sutton-Smith, M. Suzuki, H. Saito, J. C. Yeh, T. Kawano, O. Hindsgaul, P. H. Seeberger, M. Panico, S. M. Haslam, H. R. Morris, R. D. Cummings, A. Dell, M. Fukuda, Glycoconj J 2009, 26, 511–523.

[24] A. Hidalgo, A. J. Peired, M. Wild, D. Vestweber, P. S. Frenette, Immunity 2007, 26, 477–489.

[25] H. Kawashima, M. Fukuda, Ann N Y Acad Sci 2012, 1253, 112–121.

[26] a) K. Veluraja, C. J. Margulis, J Biomol Struct Dyn 2005, 23, 101–111; b) P. A. Barra, A. J. Ribeiro, M. J. Ramos, V. A. Jimenez, J. B. Alderete, P. A. Fernandes, Chem Biol Drug Des 2017, 89, 114–123.

[27] a) W. S. Somers, J. Tang, G. D. Shaw, R. T. Camphausen, Cell 2000, 103, 467–479; b) R. C. Preston, R. P. Jakob, F. P. Binder, C. P. Sager, B. Ernst, T. Maier, J Mol Cell Biol 2016, 8, 62–72.

[28] W. Thomas, J Cell Biol 2006, 174, 911–913.

[29] a) A. Barenwaldt, H. Laubli, Expert Opin Ther Targets 2019, 23, 839–853; b) C. Bull, T. Heise, G. J. Adema, T. J. Boltje, Trends Biochem Sci 2016, 41, 519–531.

[30] L. Cipolla, B. La Ferla, C. Airoldi, C. Zona, A. Orsato, N. Shaikh, L. Russo, F. Nicotra, Future Med Chem 2010, 2, 587–599.

[31] a) A. Natoni, M. S. Macauley, M. E. O’Dwyer, Front Oncol 2016, 6, 93; b) W. F. Zandberg, J. Kumarasamy, B. M. Pinto, D. J. Vocadlo, J Biol Chem 2012, 287, 40021–40030.

[32] S. J. Gamblin, S. G. Vachieri, X. Xiong, J. Zhang, S. R. Martin, J. J. Skehel, Cold Spring Harb Perspect Med 2021, 11.

