## Supporting Information for "Peracetyl *N*-cyclobutanoyl-D-mannosamine enhances expression of sialyl-Lewis X (sLeX / CD15s) and adhesion of leukocytes"

Supporting results and discussion, Supporting Figures, Supporting Movies, and Materials and Methods for

### Table of Contents

1. Supporting Results and Discussion
2. Molecular Modelling and Molecular Dynamics simulation
3. Materials and methods
4. List of reagents used in this study
5. References
6. NMR ($^1$H and $^{13}$C) spectra for **1b** and **2b**
7. Supporting Figures 1 - 14

### 1. Supporting Results and Discussion

#### 1.1. Profiling changes to global glycosylation using lectins

Apart from sLeX, the metabolic processing of ManNAc analogues could in principle affect various other types of sialoglycans on *N*- and *O*-linked glycoproteins as well as gangliosides. In order to address this, we probed global sialylation levels using sialic acid binding lectins, *viz*., *Maackia amurensis* lectin (MAL-II) and *Sambucus nigra* agglutinin (SNA) which bind, respectively, to NeuAc$\alpha$2$\rightarrow$3Gal and NeuAc$\alpha$2$\rightarrow$6Gal/GalNAc epitopes[25] (**Supporting Figure 3**). HL-60 cells incubated with vehicle (D), **1a**, **1b**, or **1d** (50 $\mu$M, 72 h) were lysed, resolved on SDS-PAGE, transferred to nitrocellulose membranes and probed with biotinylated MAL-II or biotinylated SNA followed by horse radish peroxidase conjugated avidin (HRP-avidin). MAL-II epitopes in control cells (U, D and **1d**) were found to be below detection limit with a slight increase noted upon treatment with **1a** or **1b**. MAL-II blot results were consistent with earlier observations of spontaneously induced hypo-sialylated phenotype of HL-60 cells in culture[20], [21]. By contrast, treatment of **1a** or **1b** resulted in significant reduction of SNA epitopes compared to controls (U, D, or **1d**). A comparison of lectin blots of cells (U, D, and **1d**) reveal striking differential of abundant SNA epitopes compared to low levels of MAL-II epitopes, suggesting differential endogenous activities of $\alpha$2$\rightarrow$3 *vs*. $\alpha$2$\rightarrow$6 sialyl transferases. Further evaluation of lectin binding on intact cells by flow cytometry confirmed significant gain in MAL-II binding upon treatment with **1a** or **1b**, but not in controls, compared to untreated cells (**Fig. 4C**); no significant differences were observed for SNA binding upon treatment with D, **1d**, **1a**, or **1b** compared to untreated cells (**Fig. 4D**). The increase in MAL-II with a concomitant decrease in SNA epitopes upon treatment with **1b** is in agreement with the enhanced binding observed for CSLEX1 and E-selectin-Fc since the sLeX contains the NeuAc$\alpha$2$\rightarrow$3Gal moiety.

#### 1.2. Estimation of sialic acids by periodate-resorcinol assay

In order to verify metabolic processing to sialic acids we measured both total and glycoside bound sialic acid levels using the periodate-resorcinol assay in HL-60 cells (**Supporting Figure 4**)[7, 20]. Total sialic acid levels were comparable to untreated cells for treatment with vehicle (D), the GlcNAc analogues **2a**, or **2b** (100 $\mu$M, 48 h); an increase of 1.14 to 1.3-fold was observed for treatment with **1a**, **1b**, **1c**, and **1f**. An increase of 2.0-fold and 2.3-fold was observed for **1d** and **1e**, while **1g** showed a decrease by 0.7-fold. The glycoside bound sialic acid levels were comparable to untreated cells for D, **1c**, **1f**, **2a**, and **2b**. About ~10% increase was observed for both **1d** and **1e** while both **1a** and **1b** showed a decrease of ~10%. The ratios of total sialic acids to glycoside bound sialic acids were similar to untreated cells for D, **1c**, and **1f**; higher ratios of free sialic acid were observed for **1a**, **1b**, **1d**, and **1e** while lower ratios were observed for **1g**, **2a**, and **2b**. Consistent with literature, treatment with **1d** and **1e** resulted in large increase in the metabolic flux of free sialic acid with modest gains in glycoside-bound sialic acids[7]; whereas treatment with **1a** and **1b** resulted in a gain in free sialic acids with a moderate decrease in glycoside-bound sialic acids. Our results are in agreement with the recent reports by Wittmann and coworkers wherein **1a** increased the total sialic acid levels by two folds in HEK293 cells with more than 50% being the modified *N*-cyclopropanoyl-D-neuraminic acid[21]. In the case of treatment with **1g**, **2a**, and **2b** almost all the sialic acids were glycoside bound.

Comparative analysis revealed that the *N*-cyclobutanoyl analogue **1b** resulted in increased sialic acid levels whereas the corresponding five carbon straight chain *N*-pentanoyl analogue **1g** did not show any effect, suggesting that steric effects played a prominent role; interestingly, both the *N*-cyclopropanoyl analogue **1a** and the corresponding straight chain *N*-butanoyl analogue **1f** showed similar total sialic acid levels.

### 2. Molecular Modelling and Molecular Dynamics simulation

The initial coordinates were obtained from the E-selectin– sialyl-Lewis X (sLeX) co-crystallized complex (PDB 1G1T). The cyclobutyl ring ($C_4H_7$-) was modelled onto the sLeX using PyMOL[1] by replacing the methyl group on the Neu5Ac moiety of this oligosaccharide, while maintaining the original coordinates for rest of the ligand. During preliminary structural analysis, E-selectin was found to have 10 cysteine residues and all of them were involved in the formation of five disulphide bonds, two in the lectin domain (between the residues Cys19-Cys117 and Cys90-Cys109) and three in the EGF domain (between the residues Cys122-Cys133, Cys127-Cys142, and Cys144-Cys153) (**Supporting Figure 9**). All the disulphide bonds were retained during the simulations. The $Ca^{2+}$ ion, which anchors the sLeX was not restrained during the simulations, but it was compactly held in place due to its octahedral coordination state wherein, six coordination sites were satisfied by the residue sidechains from the lectin domain of E-selectin including Glu80, Asn82, Asn83, Asn105 and Asp106 and the other two from the hydroxyl groups (O3 and O4) on the Fuc moiety of sLeX.

AMBER18 package[2] was used to carry out the MD simulations of E-selectin – sLeX and E-selectin – sLeX-Cb complexes. Force field parameters for the sLeX (wild-type) were generated from the GLYCAM webserver (www.glycam.org); for sLeX-Cb (analogue), the force field parameters for the cyclobutyl ring were generated using Antechamber[3], wherein the AM1BCC charges[4] were used. Torsional and bond parameters for the cyclobutyl ring of sLeX-Cb were curated from AMBER force field parameters[5]. TIP3P water model was used to solvate each system such that the boundary of the box was at least 14 Å away from any protein atom (thus the total number of atoms in each system was 50,903 for E-selectin Apo; 50,948 for E-selectin sLeX wild type; 50,957 for E-selectin sLeX-Cb). Counter ions ($Na^+$) were added to balance the net charges in each system. The force field ff14SB[6] was used for treating intermolecular interactions, GLYCAM06 force field[7] was used to treat sLeX and, both GLYCAM06 and Generalized Amber Force Field (GAFF)[8] were used to treat sLeX-Cb (wherein cyclobutyl ring was treated with GAFF). Particle mesh Ewald method (PME)[9] was employed for treating long range electrostatic interactions. SHAKE[10] was used for constraining all the bonds involving hydrogen. An integration time step of 2 fs was used for propagating the dynamics. Each system was initially minimized for 10,000 steps to remove any unfavourable interactions between the protein and the solvent, followed by heating to 300 K over 150 ps under normal pressure and temperature (NPT) conditions (the system was heated gradually from 0 to 300 K with an increment of 20 K in temperature after every 10ps). Subsequently, both E-selectin apo (without any ligand) and E-selectin sLeX wild complex systems were simulated for 200 ns whereas E-selectin sLeX-Cb complex system was simulated for 300 ns. Since the fluctuations in the structures were observed after 200ns for E-selectin-sLeX-Cb, the simulation time scale was extended for another 100 ns for better equilibration. All the simulations were carried out at constant temperature (300 K) and a pressure of 1.0 atm (NPT conditions), wherein the structures were stored after every 1.0 ps for analysis. In order to determine the conformational space accessed by sLeX and its analogue (sLeX-Cb) in solvent and in the absence of E-selectin, both the tetrasaccharides were simulated for 200 ns each, adding to the total simulation time of 900 ns. All the analysis were carried out using AMBER18 software, Bio3d package in R[11], and trajectories were visualised using VMD[12] and PyMOL[1].

#### Root Mean Square Deviation (RMSD)

The RMSD (using the equation given below) for C-$\alpha$ atoms with respect to the minimized structure for each MD simulation system was carried out using CPPTRAJ[13] module of AMBER18. RMSD for independent ligand simulations were calculated by superposing all atoms of the ligand with respect to its minimised structure.

$$RMSD(a,b) = \sqrt{\frac{1}{n}\sum_{i=1}^{n} |a_i - b_i|^2}$$

$$= \sqrt{\frac{1}{n}\sum_{i=1}^{n} ((a_{ix} - b_{ix})^2 + (a_{iy} - b_{iy})^2 + (a_{iz} - b_{iz})^2)}$$

where number of atoms is defined by 'n', for the target atom, '$a_i$' represents the coordinate vector whereas for reference atom it is represented by '$b_i$'. The average RMSD of 2.62 Å ± 0.73 and 2.45 Å ± 0.61 (**Supporting Figure 10A**) was observed for E-selectin from sLeX and sLeX-Cb complexes respectively. In order to delineate the effect of ligands sLeX and sLeX-Cb on E-selectin, E-Selectin in apo form (without ligands) was simulated for 200ns and average RMSD of C-$\alpha$ atoms of E-Selectin was 3.09 Å ± 1.06, which is higher than the complex simulations. All atom RMSD of ligands from the complex simulation is 2.57 Å ± 0.34 and 2.11 Å ± 0.54 for sLeX and sLeX-Cb respectively. The average all atom RMSD of sLeX wild and sLeX-Cb from ligand alone simulation is 1.87 Å ± 0.52 and 2.2 Å ± 0.59 respectively (**Supporting Figure 10B** and **Supporting Table 1**).

### Root Mean Square Fluctuation (RMSF)

The Root mean square fluctuations (RMSF), an indicator of rigid/flexible structural regions, were calculated for C-$\alpha$ atoms of E-selectin taken from the last 100ns of apo, and E-selectin complex MD simulations using respective minimized structure as the reference. Structural regions with higher flexibility in E-Selectin were mapped onto the crystal structure (PDB 1G1T).

$$RMSF(i) = \sqrt{\frac{1}{T}\sum_{i=1}^{T} |r_i(t) - \overline{r_i}|^2}$$

where 'T' is the total number of frames, '$r_i(t)$' represents the position of $i^{th}$ residues at $t^{th}$ frame and $\overline{r}_i$ gives the average position of $i^{th}$ residues with respect to frame. The RMSF observed for apo E-selectin was higher followed by E-Selectin – sLeX and E-selectin – sLeX-Cb complexes (**Supporting Figure 11A**). The structural mapping of RMSF shows higher fluctuations in the loops L4, L7, L8 and L10 (**Supporting Figure 11B**) along with both the alpha helices $\alpha1$, $\alpha2$ and beta strand $\beta5$; These structural regions are considered to be important for allosteric signalling[14].

### Characterising conformational dynamics of E-selectin

In all the three MD simulations, E-selectin was found to access multiple conformations. This change in conformation of E-selectin was characterized by opening and closing of its lectin and EGF domains which could be quantified by calculating the angle between the three points connecting the $Ca^{2+}$ ion, *C-$\alpha$* atom of Trp1 (pivot), and Cys144 residue[15]. This interdomain angle was calculated for E-selectin taken from all the snapshots of three MD simulations using CPPTRAJ. The interdomain angle $\leq$ 120° represents the bent conformation of E-selectin, whereas angle $\geq$ 145° represents the extended conformation and angle ranging from 120° – 145° represents the intermediate conformations. This cut-off for the interdomain angle was decided based on the benchmark carried out for PDB 1G1T (120°, *bent*) and PDB 4CSY (145°, *extended*). Variations in the angles as a function of time calculated for E-selectins taken from the apo, and ligand bound forms is shown in **Supporting Figure 12**. The representative structures showing conformational flexibility of E-selectin taken from apo and complex simulations are shown in **Supporting Figure 13**. As seen from **Supporting Figure 13A**, apo E-selectin takes up both bent and straight conformations, while E-selectin in complex with sLeX adapts extended conformation from early simulation time point (**Supporting Figure 13B**); by contrast, the E-selectin in complex with sLeX-Cb continues to maintain the bent conformation (**Supporting Figure 13C**).

#### Conformational dynamics of Ligands sLeX/sLeX-Cb

In order to characterise the conformational flexibility of ligands in free and in complexed form, the dihedral angles $\phi$ and $\psi$ about the glycosidic linkages were calculated (**Supporting Figure 14**). There are three glycosidic linkages in both the tetrasaccharides including Fuc$\alpha$1→3GlcNAc, Gal$\beta$1→4GlcNAc and NeuAc$\alpha$2→3Gal. The following atoms were used for dihedral angle calculation for each glycosidic linkage including, Fuc$\alpha$1→3GlcNAc: $\phi$= O5' – C1' – O3 – C3, $\psi$ = C1'- O3 – C3 – C2; Gal$\beta$1→4GlcNAc: $\phi$= O5' – C1' – O4 – C4, $\psi$ = C1'- O4 – C4 – C3 and NeuAc$\alpha$2→3Gal: $\phi$= O6' – C2' – O3 – C3, $\psi$ = C2'- O3 – C3 – C2 for each ligand[16]. Dihedral angle calculations were carried out using CPPTRAJ for the whole trajectories. While there was not much change in the torsion angles of Fuc$\alpha$1→3GlcNAc and Gal$\beta$1→4GlcNAc in the bound and unbound forms, striking differences were seen for $\phi$ and $\psi$ values of glycosidic angle of NeuAc$\alpha$2→3Gal in the bound and unbound states (Table S2 and S3). The differences in the distribution of $\phi$ and $\psi$ values of NeuAc$\alpha$2→3Gal of sLeX and sLeX-Cb complexed with E-selectin shows the widely different conformational preference of ligands in binding to E-selectin.

#### MM-GBSA – Binding energetics

As E-selectin in apo and in complex with sLeX and sLeX-Cb were found to access multiple conformations, binding energetics were calculated for the structural ensembles taken from the same complex simulations using MM/GBSA (molecular mechanics [MM] with Generalized – Born (GB and surface area (SA) model). Since, MM/GBSA method is based on many approximations and doesn't include conformational entropy[17], binding energy ($\Delta G = \Delta H - T\Delta S$) calculated from the free simulations of receptor and ligand resulted in positive $\Delta G$ values which is considered as unfavorable. However, to get the rough estimate of binding energetics of sLeX and sLeX-Cb, the snapshots taken from the respective complex simulations were considered.

Free energy for the complex formation, $\Delta G_{bind}$, is as given below:

$$\Delta G_{bind} = (\Delta G_{complex}) - (\Delta G_{Receptor}) - (\Delta G_{Ligand})$$

Originally developed by Kollman et al.,[18] wherein the components of free energy are given by

$$\Delta G_{bind} = \Delta E_{vdw} + \Delta E_{EL} + \Delta G_{GB} + \Delta G_{SURF} - T\Delta S$$

where $\Delta E_{vdw}$ and $\Delta E_{EL}$ are molecular mechanics components of bonded and non-bonded interactions (van der Waals and electrostatics). $\Delta G_{GB}$ and $\Delta G_{SURF}$ constitute the polar and non – polar energy contributions respectively to solvation free energies, where generalized Born (GB) model was used to obtain $\Delta G_{GB}$ component. $\Delta G_{SURF}$ was estimated from solvent accessible surface area (SASA). $T\Delta S$, constitutes the absolute temperature (T), multiplied by change in entropy ($\Delta S$) which was calculated using Quasi – Harmonic approximations. Binding energy calculations were done for the last 100 ns in each system (101 – 200 ns for E-selectin sLeX and 201 – 300 ns for E-selectin sLeX-Cb complexes), wherein, enthalpy (ΔH) calculations were done for four clusters (different conformations) with each cluster comprising of 25,000 frames (each cluster consists of 25 ns) whereas entropy calculations (TΔS) were done for 25 frames in all four clusters taken at an interval of 1 ns for both systems using MMPBSA.py[19] module of AMBER18 [Table S4]. The difference was seen to stem mainly from electrostatic components due to favourable interactions of carboxylate moiety of sLeX-Cb with E-selectin.

**Computational information**

Images depicting the structural insights given in the manuscript were generated using PyMOL[1], whereas the movies and the gif files from the MD simulation trajectories were generated using VMD program[12]. R program and Xmgrace program[20] (http://plasma-gate.weizmann.ac.il/Grace/) were used for plotting all the graphs.

**References**

[1] W. L. DeLano, *https://www. pymol. org* **2002**.
[2] D. A. Case, Ben-Shalom, I. Y., Brozell, S. R., Cerutti, D. S., Cheatham, T. E., Cruzeiro, V. W., Darden, T. A., Duke, R. E., Ghoreishi, D., Gilson, M. K., Gohlke, H., Goetz, A. W., Greene, D., Harris, R., Homeyer, N., Izadi, S., Kovalenko, A., Kurtzman,T., Lee, T. S., ... Kollman, P. A., University of California, **2018**.
[3] J. Wang, W. Wang, P. A. Kollman, D. A. Case, *Journal of molecular graphics and modelling* **2006**, *25*, 247-260.
[4] A. Jakalian, D. B. Jack, C. I. Bayly, *Journal of computational chemistry* **2002**, *23*, 1623-1641.
[5] W. D. Cornell, P. Cieplak, C. I. Bayly, I. R. Gould, K. M. Merz, D. M. Ferguson, D. C. Spellmeyer, T. Fox, J. W. Caldwell, P. A. Kollman, *Journal of the American Chemical Society* **1995**, *117*, 5179-5197.
[6] J. A. Maier, C. Martinez, K. Kasavajhala, L. Wickstrom, K. E. Hauser, C. Simmerling, *Journal of chemical theory and computation* **2015**, *11*, 3696-3713.
[7] K. N. Kirschner, A. B. Yongye, S. M. Tschampel, J. Gonzalez-Outeirino, C. R. Daniels, B. L. Foley, R. J. Woods, *J Comput Chem* **2008**, *29*, 622-655.
[8] J. Wang, R. M. Wolf, J. W. Caldwell, P. A. Kollman, D. A. Case, *Journal of computational chemistry* **2004**, *25*, 1157-1174.
[9] T. Darden, D. York, L. Pedersen, *The Journal of chemical physics* **1993**, *98*, 10089-10092.
[10] W. F. Van Gunsteren, H. J. C. Berendsen, *Molecular Physics* **1977**, *34*, 1311-1327.
[11] B. J. Grant, A. P. Rodrigues, K. M. ElSawy, J. A. McCammon, L. S. Caves, *Bioinformatics* **2006**, *22*, 2695-2696.
[12] W. Humphrey, A. Dalke, K. Schulten, *Journal of molecular graphics* **1996**, *14*, 33-38.
[13] D. R. Roe, T. E. Cheatham, 3rd, *J Chem Theory Comput* **2013**, *9*, 3084-3095.
[14] T. A. Springer, *Proceedings of the National Academy of Sciences* **2009**, *106*, 91-96.
[15] R. C. Preston, R. P. Jakob, F. P. Binder, C. P. Sager, B. Ernst, T. Maier, *J Mol Cell Biol* **2016**, *8*, 62-72.
[16] R. M. Cooke, R. S. Hale, S. G. Lister, G. Shah, M. P. Weir, *Biochemistry* **1994**, *33*, 10591-10596.
[17] S. Genheden, U. Ryde, *Expert opinion on drug discovery* **2015**, *10*, 449-461.
[18] J. Srinivasan, T. E. Cheatham, P. Cieplak, P. A. Kollman, D. A. Case, *Journal of the American Chemical Society* **1998**, *120*, 9401-9409.
[19] B. R. Miller, 3rd, T. D. McGee, Jr., J. M. Swails, N. Homeyer, H. Gohlke, A. E. Roitberg, *J Chem Theory Comput* **2012**, *8*, 3314-3321.
[20] P. J. Turner, *Center for Coastal and Land-Margin Research, Oregon Graduate Institute of Science and Technology, Beaverton, OR* **2005**, *2*.

### 3. Materials and methods

#### 3.1. Synthesis of HexNAc analogues and their characterization

**General information**: All chemicals used were of analytical grade and were used as such unless mentioned otherwise. Hexosamine starting materials, namely, D-Mannosamine hydrochloride, D-glucosamine hydrochloride, D-galactosamine hydrochloride, reagents, and solvents were purchased commercially. Compounds were purified by flash column chromatography using silica gel 60 (230-400 mesh) manually. Solvents used for chromatography were either analytical grade or distilled prior to use. Solvent evaporations were performed using rotary evaporator. Thin layer chromatography (TLC) was performed using fluorescent silica gel glass plates (Analtech / Miles Scientific) and visualised using handheld UV lamp and mostain. NMR ($^1$H and $^{13}$C) was recorded on a Bruker 300 MHz spectrometer using chloroform-*d*, methanol-$d_4$, dimethyl sulfoxide-$d_6$, and deuterium oxide as solvents with tetramethylsilane (TMS) as internal reference. Mass spectrometry was performed using high resolution ESI-MS (Thermo Orbitrap Velos) or MALDI-TOF-TOF (AB Sciex 4800) instruments. For MALDI, super-DHB was used as matrix.

The *N*-acyl-D-hexosamine analogues used in this study were synthesized following previously reported procedures and characterized using TLC, NMR ($^1$H and $^{13}$C), and Mass spectrometry. Compounds **1a**[1], **1c**[2-4], **1d**[4, 5], **1e**[4], **1f**[3-5], **1g**[4], **2a**[1], and **3**[6] are previously reported and compounds **1b** and **2b** are novel compounds. Detailed characterizations are given below for all the compounds since we employed single anomers, i. e., the α-anomers for ManNAc series (**1a-1g**) and the β-anomers for both GlcNAc series (**2a** and **2b**) and Ac$_4$GalNAz (**3**). The characterization data were in agreement with published values where available.

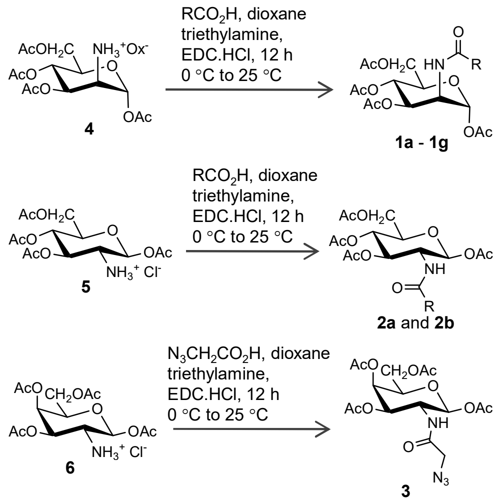

**Supporting Scheme 1. Synthesis of HexNAc analogues**. Peracetylated hexosamine salts, **4**, **5**, and **6**, were synthesized according to reported procedures and subjected to carbodiimide mediated coupling with various carboxylic acids to obtain the HexNAc analogues **1a** – **1g**, **2a**, **2b**, and **3** which were employed in this study.

Selectively protected hexosamine intermediates, namely, 1,3,4,6-tetra-*O*-acetyl-2-amino-2-deoxy-α-D-mannosamine oxalic acid salt (**4**)[7], 1,3,4,6-tetra-*O*-acetyl-2-amino-2-deoxy-β-D-glucosamine hydrochloride salt (**5**)[8], and 1,3,4,6-tetra-*O*-acetyl-2-amino-2-deoxy-β-D-galactosamine hydrochloride salt (**6**)[9] were synthesized according to the reported procedures. The respective peracetylated hexosamine salts were subjected to carbodiimide mediated coupling with various carboxylic acids to obtain the HexNAc analogues (Supporting Scheme S1).

**1,3,4,6-Tetra-*O*-acetyl-2-cyclopropanoylamino-2-deoxy-α-D-mannopyranose** ($Ac_4$Man-NCp, **1a**): To a stirred suspension of $Ac_4ManNH_3^+Ox^-$ (**4**) (1.0 g, 2.28 mmol), cyclopropanoic acid (0.59 mg, 6.84 mmol), and EDC. HCl (0.874 g, 4.56 mmol) in dioxane (20 mL), at 25 °C, was added triethylamine (6.0 mmol). After 16 h, reaction mixture was concentrated, residue was dissolved in dichloromethane (100 mL) and washed with aq. $NaHCO_3$ (5 % w/v). Organic layer was separated, dried (anh. $Na_2SO_4$), filtered, concentrated, and the residue was purified by silica gel column chromatography using hexanes / ethyl acetate as eluant to obtain pure **1a** (0.3 g, 31 %). The NMR data was found to be identical to those reported in the literature[1].

$^1$H NMR (300 MHz, $CDCl_3$): δ 6.05 (d, *J* = 1.8 Hz, 1H), 5.93 (d, *J* = 9.0 Hz, 1H), 5.32 (dd, *J* = 10.2 Hz,4.2 Hz, 1H), 5.21 (dd, *J* = 9.9 Hz, 9.8 Hz 1H), 4.66 (ddd, *J* = 9.0, 4.2, 1.8 Hz, 1H), 4.29 (dd, *J* = 12.6Hz, 4.8 Hz, 1H), 4.06 (dd, 12.0 Hz, 2.7 Hz, 1H), 4.05 (m, 1H), 2.17 (s, 3H), 2.10 (s, 3H), 2.07 (s, 3H),1.98 (s, 3H), 1.43 (m, 1H), 0.97 (m, 2H), 0.83 (m, 2H); $^{13}$C NMR (75 MHz, $CDCl_3$): δ 173.8, 170.5,170.0, 169.8, 168.1, 91.8, 70.10, 68.9, 65.6, 62.1, 49.3, 20.8, 20.74, 20.67 (2C), 14.8, 8.01, 7.96; ESI-MS m/z Calcd for $[C_{18}H_{25}NO_{10}+ Na]^+$ 438.1370; Found 438.1370.

**1,3,4,6-Tetra-*O*-acetyl-2-cyclobutanoylamino-2-deoxy-α-D-mannopyranose** ($Ac_4$Man-NCb, **1b**): To a stirred suspension of EDC.HCl (0.44 g, 2.3 mmol, 2.0 equiv.) in dioxane (10 mL) was added cyclobutanoic acid (0.22 mL, 2.3 mmol, 2.0 equiv.) at 25 °C, followed by $Ac_4ManNH_3^+$ $Ox^-$ (**4**) (0.504 g, 1.15 mmol, 1.0 equiv.) and triethylamine (2.4 mL, 17 mmol, 15 equiv.). Instant formation of colorless precipitate was noticed and the reaction was allowed to stir overnight. The reaction mixtures concentrated, the residue was dissolved in dichloromethane (50 mL) and washed with satd. aq. $NaHCO_3$ solution (50 mL). The organic layer was dried (anh. $Na_2SO_4$), filtered, concentrated, and the residue was purified by silica gel column chromatography to obtain pure **1b** (0.23 g, 46 %).

$^1$H NMR (300 MHz, $CDCl_3$): δ 6.01 (d, *J* = 1.8 Hz, 1H) ,5.53 (d, *J* = 9.3 Hz, 1H), 5.32 (dd, *J* = 10.2, 4.5 Hz, 1H), 5.12 (dd, *J* = 10.2 Hz, 1H), 4.65 (ddd, *J* = 9.3, 4.5, 1.8 Hz, 1H), 4.25 (dd, *J* = 4.5, 12.3 Hz, 1H), 4.05 (dd, *J* = 2.4, 12.3 Hz, 1H), 4.03 (ddd, *J* = 2.4, 4.2, 8.4 Hz, 1H), 3.06 (q, *J* = 8.4 Hz, 1H), 2.27 (m, 4H), 2.17 (s, 3H), 2.09 (s, 3H), 2.06 (s, 3H), 2.00 (s, 3H),1.96 (m, 2H); $^{13}$C NMR (75 MHz, $CDCl_3$): δ 175.0, 170.5, 170.0 169.7, 168.1, 91.8, 70.1, 68.9, 65.4, 62.0, 49.0, 39.7, 25.32, 25.3, 20.97, 20.8, 20.7, 20.6, 18.2; NMR spectra of **1b** are provided below. ESI-MS m/z Calcd for $[C_{19}H_{27}NO_{10}+ Na]^+$ 452.1527; Found 452.1526.

**1,3,4,6-Tetra-*O*-acetyl-2-azidoacetylamino-2-deoxy-α-D-mannopyranose** ($Ac_4$ManNAz, **1c**): Synthesis and characterization of $Ac_4$ManNAz (**1c**) has been reported by us previously and the data was found to agree with earlier reports.[2, 3]

**1,3,4,6-Tetra-*O*-acetyl-2-acetylamino-2-deoxy-α/β-D-mannopyranose** ($Ac_4$ManNAc, **1d**): Synthesis of **1d** was performed as reported previously and the data was found to agree with earlier reports.[4, 5]

**1,3,4,6-Tetra-*O*-acetyl-2-(1-propanoyl)amino-2-deoxy-α-D-mannopyranose** ($Ac_4$Man-NProp, **1e**): To a solution of $Ac_4ManNH_3^+$ $Ox^-$ (**4**)[7] (0.5 g, 1.14 mmol) in DMF (15 mL), cooled in an ice-water bath, was added propanoic anhydride (0.325 mL, 2.52 mmol, 2.2 equiv.), dropwise with stirring, followed by triethylamine (0.48 mL, 3.43 mmol, 3.0 equiv.) and allowed to warm up to room temperature. After 12 h, the reaction mixture was concentrated, residue was dissolved in dichloromethane (50 mL), and washed with aq. $NaHCO_3$ (2.0 % w/v) (50 mL). The aqueous layer was washed with dichloromethane (25 mL × 2). The organic layers were combined, dried (anh. $Na_2SO_4$), filtered, and concentrated. The residue was purified by silica gel column chromatography to obtain pure **1e** (0.32 g, 69 %). The NMR data was found to be in agreement with reported values.[4, 5]

$^1$H NMR (300 MHz, $CDCl_3$): δ 6.02 (d, *J* = 1.8 Hz, 1H), 5.70 (d, *J* = 9.3 Hz, 1H), 5.33 (dd, *J* = 4.5, 10.0 Hz, 1H), 5.16 (dd, *J* = 10.0 Hz, 1H), 4.65 (ddd, *J* = 1.8, 4.2, 9.0 Hz, 1H), 4.27 (dd, *J* = 5.1, 12.0, Hz, 1H), 4.08-4.01 (m, 2H), 2.29 (q, *J* = 7.2 Hz, 2H), 2.18 (s, 3H), 2.10 (s, 3H), 2.06 (s, 3H), 2.00 (s, 3H), 1.18 (t, *J* = 7.0 Hz, 3H); $^{13}$C NMR (75 MHz, $CDCl_3$): δ 173.9, 170.5, 170.0, 169.7, 168.2, 91.8, 70.1, 68.9, 65.5, 62.1, 49.1, 29.6, 20.8, 20.7 (2C), 20.6, 9.7; ESI-MS (m/z): Calcd for $C_{17}H_{25}NO_{10}Na$ [M+ Na]$^+$,426.1376; found,426.1364.

**1,3,4,6-Tetra-*O*-acetyl-2-(1-butanoyl)amino-2-deoxy-α-D-mannopyranose** ($Ac_4$ManNBut, **1f**): To a solution of $Ac_4ManNH_3^+$ $Ox^-$ (**4**)[7] (0.5 g, 1.14 mmol) in DMF (15 mL), cooled in an ice-water bath, was added butanoic anhydride (0.41 mL, 2.52 mmol, 2.2 equiv.), dropwise with stirring, followed by triethylamine (0.48 mL, 3.43 mmol, 3.0 equiv.) and allowed to warm up to room temperature. After 12 h, the reaction mixture was concentrated, residue was dissolved in dichloromethane (50 mL), and washed with aq. $NaHCO_3$ (2.0 % w/v) (50 mL). The aqueous layer was washed with dichloromethane (25 mL × 2). The organic layers were combined, dried (anh. $Na_2SO_4$), filtered, and concentrated. The residue was purified by silica gel column chromatography to obtain pure **1f** (0.35 g, 73 %). The NMR data was found to be identical to reported values in the literature.[3-5]

$^1$H NMR (300 MHz, $CDCl_3$): δ 6.02 (d, *J* = 1.8 Hz, 1H), 5.71 (d, *J* = 9.0 Hz, 1H), 5.33 (dd, *J* = 4.5, 10.2 Hz, 1H), 5.17 (dd, *J* = 10.2 Hz, 1H), 4.66 (ddd, *J* = 1.8, 4.2, 9.0 Hz, 1H), 4.28 (dd, *J* = 5.1, 12.6 Hz, 1H), 4.09-4.01 (m, 2H), 2.24 (t, *J* = 7.5 Hz, 2H), 2.18 (s, 3H), 2.10 (s, 3H), 2.06 (s, 3H), 1.99 (s, 3H), 1.68 (m, 2H), 0.99 (t, *J* = 7.2 Hz, 3H); $^{13}$C NMR (75 MHz, $CDCl_3$): δ 173.0, 170.5, 170.0, 169.7, 169.2, 91.8, 70.1, 68.9, 65.5, 62.0, 49.0, 38.5, 20.8, 20.7, 20.6, 20.4, 19.1, 13.6; ESI-MS (m/z): Calcd for $C_{18}H_{27}NO_{10}Na$ [M+ Na]$^+$, 440.1533; found, 440.1520.

**1,3,4,6-Tetra-*O*-acetyl-2-(1-pentanoyl)amino-2-deoxy-α-D-mannopyranose** ($Ac_4$ManNPent, **1g**): To a solution of $Ac_4ManNH_3^+$ $Ox^-$ (**4**)[7] (0.4 g, 0.92 mmol) in DMF (10 mL), cooled in an ice-water bath, was added valeric anhydride (0.4 mL, 2.0 mmol, 2.2 equiv.), dropwise with stirring, followed by triethylamine (0.38 mL, 2.67 mmol, 2.9 equiv.) and allowed to warm up to room temperature. After 12 h, the reaction mixture was concentrated, residue was dissolved in dichloromethane (50 mL), and washed with aq. $NaHCO_3$ (2.0 % w/v) (50 mL). The aqueous layer was washed with dichloromethane (25 mL × 2). The organic layers were combined, dried (anh. $Na_2SO_4$), filtered, and concentrated. The residue was purified by silica gel column chromatography to obtain pure **1g** (0.17 g, 43 %). The NMR data was found to be in agreement with reported values.[4, 5]

$^1$H NMR (300 MHz, $CDCl_3$): δ 6.00 (d, *J* = 1.8 Hz, 1H), 5.78 (d, *J* = 9.3 Hz, 1H), 5.31 (dd, *J* =10.2 Hz, 1H), 5.16 (dd, *J* = 9.9, 10.2 Hz, 1H), 4.65 (ddd, *J* = 1.8, 4.2, 9.0 Hz, 1H), 4.27 (dd, *J* = 5.1, 12.6 Hz, 1H), 4.07-4.00 (m, 2H), 2.25 (t, *J* = 7.5 Hz, 2H), 2.17 (s, 3H), 2.09 (s, 3H), 2.05 (s, 3H),1.98 (s, 3H), 1.63 (m, 2H), 1.37 (m, 2H), 0.93 (t, *J* = 7.2 Hz, 3H); $^{13}$C NMR (75 MHz, $CDCl_3$): δ 173.1, 170.5, 170.0, 169.6, 168.1, 91.8, 70.1, 68.9, 65.4, 62.0, 49.1, 36.3, 27.7, 22.2, 20.8, 20.7 (2C), 20.6, 13.7; ESI-MS (m/z): Calcd. For $C_{19}H_{29}NO_{10}Na$ [M+ Na]$^+$, 454.1689; found, 454.1671.

General procedure for the synthesis of **2a** and **2b**: To a stirred suspension of a mixture of $Ac_4GlcNH_3^+ Cl^-$ (**5**) (1.0 mmol), either cyclopropanoic acid or cyclobutanoic acid (3.0 mmol), and EDC. HCl (2.0 mmol) in DMF (20 mL), at 25°C, was added triethylamine (6.0 mmol). After 16 h, the reaction mixture was concentrated, residue was dissolved in dichloromethane (100 mL), and washed with aq. $NaHCO_3$ (5 % w/v). Organic layer was separated, dried (anh. $Na_2SO_4$), filtered, concentrated, and the residue was purified by silica gel column chromatography using hexanes / ethyl acetate as eluant.

**1,3,4,6-Tetra-*O*-acetyl-2-cyclopropanoylamino-2-deoxy-β-D-glucopyranose** ($Ac_4$GlcNCp, **2a**): The NMR data was found to be identical to reported values.[1] 630 mg (58 %). $^1$H NMR (300 MHz, $CDCl_3$): δ 5.92 (d, *J* = 9.0 Hz, 1H), 5.69 (d, *J* = 9.0 Hz, 1H), 5.17 (dd, *J* = 9.6 Hz, 9.6 Hz,1H), 5.14 (dd, *J* = 9.3 Hz, 9.3 Hz,1H), 4.37 – 4.26 (m, 1H), 4.26 (dd, *J* = 12.3 Hz, 4.2 Hz, 1H), 4.11 (dd, *J* = 12.3 Hz, 4.2 Hz, 1H), 3.81 (ddd, *J* = 8.7 Hz, 4.8 Hz, 2.4 Hz, 1H), 2.10 (s, 3H), 2.08 (s, 3H), 2.03 (s, 6H), 1.27 (m, 1H), 0.93 – 0.88 (m, 2H), 0.77 – 0.70 (m, 2H). $^{13}$C NMR (75 MHz, $CDCl_3$) δ 173.8, 171.2, 170.7, 169.6, 169.3, 92.7, 72.9, 72.6, 67.8, 61.7, 52.9, 20.8, 20.7, 20.6, 20.5, 14.6, 7.44, 7.40. ESI-MS m/z Calcd for $[C_{18}H_{25}NO_{10}+ Na]^+$ 438.1370; Found 438.1368.

**1,3,4,6-Tetra-*O*-acetyl-2-cyclobutanoylamino-2-deoxy-β-D-glucopyranose** ($Ac_4$GlcNCb, **2b**): 1.11 g (92 %); $^1$H NMR (300 MHz, $CDCl_3$) δ 5.68 (d, *J* = 8.7 Hz, 1H), 5.53 (d, *J* = 9.6 Hz, 1H), 5.18 (dd, *J* = 9.3 Hz, 9.3 Hz, 1H), 5.13 (dd, *J* = 9.3 Hz, 9.3 Hz, 1H), 4.37 – 4.26 (m, 1H), 4.25 (dd, *J* = 12.6 Hz, 4.8 Hz, 1H), 4.12 (dd, *J* = 12.6 Hz, 2.4 Hz, 1H), 3.80 (ddd, *J* = 8.7 Hz, 4.2 Hz, 2.7 Hz, 1H), 2.90 (q, *J* = 8.4 Hz, 1H), 2.20 – 1.70 (m, 6H), 2.09 (s, 3H), 2.08 (s, 3H), 2.03 (s, 3H), 2.02 (s, 3H); $^{13}$C NMR (75 MHz, $CDCl_3$) δ 175.0, 171.2, 170.7, 169.5, 169.3, 92.7, 72.9, 72.6, 67.8, 61.7, 52.7, 39.8, 25.2 (2C), 20.8, 20.7, 20.62, 20.57, 18.11; NMR spectra of **2b** are provided below. ESI-MS m/z Calcd for $[C_{19}H_{27}NO_{10}+ Na]^+$ 452.1527; Found 452.1525.

**1,3,4,6-Tetra-*O*-acetyl-2-azidoacetylamino-2-deoxy-β-D-galactopyranose** ($Ac_4$GalNAz, **3**): Synthesis of **3** was performed as reported previously[10] starting from **6** and the characterization data was found to agree with reported values.[6]

### 3.2. Experiments in cell cultures in vitro

Jurkat cells (human T cell leukemia) were obtained as a kind gift from Dr. Ayub Qadri, National Institute of Immunology, New Delhi, and HL-60 cells (human acute myeloid leukemia) were purchased from European collection of authenticated cell cultures (ECACC, UK). Cells were cultured in flasks or petri dishes, under sterile conditions, in RPMI 1640 supplemented with 10% fetal bovine serum (FBS) and 1% P/S (50 U/mL of penicillin and 0.05 mg/mL of streptomycin) at 37 °C in a humidified incubator maintaining 5 % carbon dioxide. Cell culture work was performed inside a Baker SterilGard biosafety cabinet. Cells were counted using a Beckman Coulter Z2 particle counter in 0.45 μm filtered phosphate buffered saline (PBS) or haemocytometer.

Stock solutions of peracetylated HexNAc analogues **1a** – **1g**, **2a**, **2b**, and **3** (50 mM) were prepared in DMSO (**D**) and were filtered through 0.22 μm PTFE syringe filter. Stock solutions were added directly to cell culture keeping the total volume of DMSO to a maximum of 0.25 % v/v. All the experiments were initiated using actively dividing log phase cells. Cells were seeded at a density of $3.0 \times 10^5$ cells/mL for all the experiments, unless mentioned otherwise. Dibenzocyclooctyne-Cy5 (DBCO-Cy5; stock solution of 10 mM in DMSO) was employed for estimation of NeuAz expression using strain promoted azide-alkyne cycloaddition (SPAAC). Propidium iodide (PI) (stock solution of 1.0 mg/mL in water) was used to exclude non-viable cells. Jurkat cells (untreated, U), without treatment of vehicle, were employed as additional control in all the experiments. Cell counts were taken directly from the media, after gentle pipetting to single cell suspension, at the time of harvest in order to obtain a measure of cytotoxicity due to the analogues. FACS staining buffer (FSB), consisting of 1.0 % BSA w/v and 0.05 % sodium azide w/v in PBS, was used for flow cytometry experiments.

#### 3.2.1. Estimation of NeuAz expression by flow cytometry

Jurkat cells ($3.0 \times 10^5$ cells/mL; 2.5 mL per well in 6-well plates) in complete medium were treated with either vehicle (**D**) or $Ac_4$ManNAz (**1c**); Untreated Jurkat cells, without addition of vehicle, were also included as additional controls. Cells were harvested at 48 h, washed thrice with PBS (3 × 0.5 mL) (3000×*g*, 2.0 min). Prior to the last wash cells were counted and were aliquoted to $1.0 \times 10^6$ cells per sample in 1.5 mL microcentrifuge tubes and centrifuged. The cells were gently resuspended in PBS (50 μL) and treated with DBCO-Cy5 (5.0 μM) and incubated at 37 °C (on a Torrey-Pines dry heating bath with mild shaking). After one hour, the cells were washed using FSB (0.5 mL, 3000×*g*, 2.0 min, room temperature), re-suspended in PBS (250 μL), treated with PI (2.5 μg/mL), and analyzed using BD FACSCanto and FACSDiva Version 6.1.3. The cells were gated for PI-negative population for 10,000 events. Two replicate samples were prepared for each condition and each sample was counted twice.

For the optimization of DBCO-Cy5 concentration for SPAAC, Jurkat cells cultured with D or **1c** (50 μM) for 24 h were harvested, washed, and aliquoted in to $5.0 \times 10^5$ cells per sample in PBS (50 μL). The cells were treated with DBCO-Cy5 at 0, 5, 10, 20, and 40 μM concentrations at 37 °C. After 1.0 h, cells were washed in FSB (3 × 0.5 mL at 3000×*g* for 2.0 min), resuspended in PBS (250 μL), treated with PI (2.5 μg/mL), and analyzed by BD FACSVerse and FlowJo V10. The cells were gated for PI-negative population for 10,000 events. Two replicate samples were prepared for each condition and each sample was counted twice.

For the optimization of incubation time for SPAAC using DBCO-Cy5, Jurkat cells ($3.0 \times 10^5$ cells/mL in 5.0 mL cultured in 60 mm diameter petri dishes) incubated with either **D** or **1c** (50 μM) for 24 h were harvested and washed with PBS (3 × 0.5 mL at 3000×*g* for 2.0 min).

Cells were aliquoted at $5.0 \times 10^5$ cells per sample and treated with DBCO-Cy5 (20 μM) at 37 °C. Reactions were stopped at various time points (0 – 120 min) by washing with FSB ($3 \times 0.5$ mL), resuspended in PBS (250 μL), and kept on ice-bath. Cells were treated with PI and analyzed by BD FACSVerse and FlowJo V10 as mentioned above. Based on the results of optimization experiments, we chose incubation time of 30 min at 37 °C for $5.0 \times 10^5$ cells (pre-incubated with **1c** for 24 h) with 20 μM of DBCO-Cy5 in 50 μL total volume for experiments with ManNAc analogues.

*Estimation of NeuAz expression by **1c** in the presence of peracetylated ManNAc analogues*: Jurkat cells ($3.0 \times 10^5$ cells/mL in 3.0 mL per well in 6-well plates) were incubated with (i) D, (ii) **1a**, **1b**, **1c**, or **1d** alone (50 μM) or (iii) **1a**, **1b**, or **1d** (50 μM) in combination with **1c** (50 μM). After 24 h, cells were harvested, washed using FSB, aliquoted to $5.0 \times 10^5$ cells per sample, and processed as mentioned above for estimation of NeuAz expression using DBCO-Cy5 (20 μM). Experiments using HL-60 cells were performed using the same procedure.

*Estimation of NeuAz expression upon simultaneous and delayed addition of **1c***: In one condition, Jurkat cells were incubated with **D**, **1a**, or **1b** (50 μM) along with **1c** (50 μM). In another condition, Jurkat cells were first incubated with **D**, **1a**, or **1b** (50 μM) alone followed by the addition of **1c** (50 μM) at 12 h. After a total of 36 h, cells were harvested, washed with PBS ($3 \times 0.5$ mL), aliquoted to $5.0 \times 10^5$ cells per sample in 250 □L, reacted with DBCO-Cy5 (20 μM), washed, treated with PI, and analyzed by flow cytometry as mentioned above.

*Effect of GlcNAc and ManNAc analogues on NeuAz expression*: Jurkat cells were treated with **1a**, **1b**, **2a**, or **2b** (10, 25, and 50 μM for each analogue) in combination with **1c** (50 □M). Cells treated with **1c** (50 μM) alone were considered as a reference. After 24 h, cells were harvested, processed, labelled with DBCO-Cy5 (20 μM), and analysed by flow cytometry as mentioned above.

*Effect of ManNAc analogues on NeuAz and GalNAz expression*: Jurkat cells were incubated under various conditions, namely, (i) **D**, **1c**, **3**, **1a**, or **1b** alone, (ii) **1a** or **1b** in combination with **1c**, (iii) **1a** or **1b** in combination with **3**. After 24 h, cells were harvested, washed, labelled using DBCO-Cy5 (20 μM), processed, and analysed by flow cytometry as mentioned above.

#### 3.2.2. Estimation of sialoglycan epitopes on the cell surface by flow cytometry

*Effect of ManNAc analogues on the expression of sialyl-Lewis-X and Lewis-X*: HL-60 cells ($3.0 \times 10^5$ cells/mL in 2.0 mL complete medium in 6 well plate) were treated with D (vehicle), **1a**, **1b**, **1c**, **1d**, **1e**, **1f**, or **1g** (50 μM), along with untreated (U) control. After 24 h, cells were harvested and washed with PBS ($3 \times 0.5$ mL, 3000×*g*, 3.0 min at room temperature). Prior to last wash, cells were counted using Z2 coulter counter and aliquoted at $2.5 \times 10^5$ cells per sample and stained with either mouse anti-human CD15s (CSLEX1) (1.25 μg/mL) or mouse anti-human CD15 (HI98) (2.5 μg/mL) in HEPES /$CaCl_2$ buffer (30 mM HEPES, 110 mM NaCl, 10 mM KCl, 2 mM $MgCl_2$, 10 mM glucose, 1.5 mM $CaCl_2$ containing 0.1% bovine serum albumin, pH 7.3) (total volume of 100 μL) for 30 min at 4 °C. Cells were washed with HEPES/$CaCl_2$ buffer ($2 \times 0.5$ mL) and stained with AlexaFluor488-conjugated anti-mouse IgM (2.5 μg/mL) (for both CD15s and CD15) in 100 μL HEPES/$CaCl_2$ for 30 min at 4 °C. Cells were washed with HEPES/$CaCl_2$ buffer ($2 \times 0.5$ mL), resuspended in PBS (0.2 mL), treated with PI (2.5 μg/ml) and analysed by BD FACSVerse. Samples treated with (i) isotype antibodies followed by secondary antibody and (ii) secondary antibody alone were prepared as controls. Single Cells were gated for PI-negative population for 10,000 events and analysed by flow cytometry. Two replicate samples were prepared for each condition and each sample was counted twice. Three independent experiments were performed. Data was analysed using FlowJo V10. For plotting graph and calculation of significant probability '*p*' values, two-way ANOVA with Dunnett's multiple comparison tests was employed in GraphPad Prism 8.

*Time course of CD15s expression*: HL-60 cells (3.0 × $10^5$ cells/mL in 2.5 mL complete medium in 6-well plates) were incubated with **D**, **1a**, **1b**, or **1d** (50 μM). At each time point, from day 1 to day 6, the cells were harvested from the entire contents of the well, washed, and aliquoted to 5.0 × $10^5$ cells per sample, and immunostained using anti-CD15s (CSLEX1) antibody as mentioned above. On day 3, 4, and 5 complete media (1.0 mL) was added to each remaining well in order to compensate for media exhaustion. No contents were removed from the well during the period.

*Comparison of mouse and human E-selectin binding*: HL-60 cells were harvested from actively growing cultures and washed with PBS (3 × 0.5 mL). Prior to the last wash, cells were counted and aliquoted in to 5.0 × $10^5$ cells per sample. Cells were incubated with either human E-selectin-Fc chimera protein (hCD62E-Fc; 2.5 μg/mL) or mouse E-selectin-Fc chimera protein (mCD62E-Fc; 2.5 μg/mL) in PBS (50 μL) for 45 min at 4 °C. Cells were then washed with PBS (2 × 0.5 mL), stained with PE-conjugated $F(ab')_2$ goat anti-human Fc (2.5 μg/mL) in PBS (50 μL) for 45 min at 4 °C. Cells were then washed with PBS (2 × 0.5 mL), resuspended in 0.85% saline (0.2 mL) containing Sytox Red (dilution of 1:1000; stock solution of 5.0 μM in DMSO), and analyzed by flow cytometry using BD FACSVerse. Cells were gated on Sytox Red negative population and 10,000 events were counted within the gate. Two replicate samples were analyzed for each condition and each sample was counted twice. Data were analyzed using FlowJo V10.

*Optimization of concentration of mouse E-selectin-Fc chimera protein*: HL-60 cells were harvested from actively growing cultures and washed with PBS (3 × 0.5 mL). Cells were counted, aliquoted in to 5 × $10^5$ cells per sample, treated with mouse E-selectin-Fc chimera protein (mCD62E-Fc) at 0, 2.5, 5.0, 10, and 20 μg/mL concentrations in PBS (50 μL) and incubated at 4 °C. After 1 h, cells were washed with PBS (2 × 0.5 mL) and stained with PE-conjugated $F(ab')_2$ goat anti-human Fc in PBS (50 μL) at 4 °C. After 1 h, cells were washed with PBS (2 × 0.5 mL) and resuspended in 0.85 % saline (0.2 mL) containing Sytox Red (dilution 1: 1000; stock solution of 5.0 μM in DMSO). Two replicate samples were analyzed for each condition and each sample was counted twice by flow cytometry as mentioned above.

*Effect of HexNAc analogues on the expression of E-selectin ligands*: HL-60 cells (3.0 × $10^5$ cells/mL in 2.5 mL complete medium in six-well plates) were incubated with D or 1a-1g (50 μM), along with untreated controls. After 24 h, cells were harvested and washed with PBS (3 × 0.5 mL). Prior to the last wash, cells were counted and aliquoted into 2.5 × $10^5$ cells per sample. Cells were stained with mCD62E-Fc chimera protein (2.5 μg/mL) in HEPES/$CaCl_2$ (100 μL) at 4 °C. After 30 min, cells were washed with HEPES/$CaCl_2$ (2 × 0.5 mL) and stained with PE-conjugated F(ab')2 goat anti-human Fc in HEPES/$CaCl_2$ (100 μL) at 4 °C. After 30 min, cells were washed with HEPES/$CaCl_2$ (2 × 0.5 mL) and resuspended in HEPES/$CaCl_2$ (0.2 mL) containing Sytox Red (dilution 1:1000; stock concentration of 5.0 μM). Additionally, control samples of cells treated with only the secondary antibody were prepared. Two replicate samples were analyzed for each condition and each sample was counted twice by flow cytometry as mentioned above. Two independent experiments were performed.

*Effect of HexNAc analogues on the expression of cutaneous lymphocyte antigen (CLA / HECA452) epitopes*: HL-60 cells (3.0 × $10^5$ cells/mL in 2.5 mL complete medium in 6-well plates) were treated with **D** or **1a-1g** (50 μM), including untreated controls. After 24 h cells were harvested, washed in PBS (3 × 0.5 mL), aliquoted to 2.5 × $10^5$ cells per sample, and incubated with rat anti-human CLA (HECA452) antibody (1.25 μg/mL) in HEPES/$CaCl_2$ (100 μL) at 4 °C. After 30 min, cells were washed with HEPES/$CaCl_2$ (2 × 0.5 mL) and stained with AlexaFluor488- conjugated anti-rat IgM (2.5 μg/mL) in HEPES/$CaCl_2$ (100 μL) at 4 °C. After 30 min, cells were washed with HEPES/$CaCl_2$ (2 × 0.5 mL), resuspended in HEPES/$CaCl_2$ (0.2 mL) and treated with PI (2.5 μg/mL). Additionally control samples of cells treated with (i) isotype antibody followed by secondary antibody and (ii) only secondary antibody were prepared. Cells were gated on PI-negative population and analyzed by flow cytometry as given above. Two replicate samples were analyzed for each condition and each sample was counted twice by flow cytometry as mentioned above. Two independent biological experiments were performed.

*Effect of ManNAc analogues on MAL-II and SNA epitopes*: HL-60 cells (3.0 × $10^5$ cells/mL in 2.5 mL complete medium in six-well plates) were treated with **D**, **1a**, **1b**, or **1d** (50 μM), along with untreated controls. After 24 h, cells were harvested, washed with PBS (3 × 0.5 mL), aliquoted in to 5.0 × $10^5$ cells per sample, and incubated with either biotinylated *Maackia amurensis* lectin-II (bMAL-II; 10 μg/mL) or biotinylated *Sambucus nigra* agglutinin (bSNA; 5.0 μg/mL) in ice-cold PBS (50 μL) at 4 C. After 30 min, cells were washed with ice-cold PBS (2 × 0.5 mL), followed by staining with FITC-conjugated avidin (dilution 1:250; stock solution of 1.0 mg/mL) in PBS (50 μL). After 30 min cells were washed with ice-cold PBS (2 × 0.5 mL), resuspended in PBS (0.2 mL), and treated with PI (2.5 μg/mL). Cells were gated on PI-negative population and analyzed by flow cytometry as given above. Two replicate samples were analyzed for each condition and each sample was counted twice by flow cytometry as mentioned above.

#### 3.2.3. Western and avidin (far-western) blotting experiments

*Effect of ManNAc analogues on expression of CD15s epitopes*: HL-60 cells (3.0 × $10^5$ cells/mL in 10 mL complete medium in 10 cm diameter petri dishes) were incubated with **D**, **1a**, **1b**, **1d**, **1e**, **1f**, or **1g** (50 μM), along with untreated (U) controls. After 72 h, cells were harvested, washed with PBS (3 × 5.0 mL), and lysed in RIPA buffer ($10^7$ cells in 100 μL), containing protease inhibitor cocktail (PIC, 1:100). Protein concentration of the soluble fraction was estimated using the Bradford assay. Proteins from total lysates were resolved by 7.5 % SDS-PAGE and blotted onto nitrocellulose membrane (constant current of 250 mA for 3.0 h at 4 °C). Membranes were blocked using non-fat milk (NFM) (5.0 % w/v) in PBS and incubated with primary antibodies (5.0 mL PBS containing 5.0% NFM) overnight at 4 °C. Membranes were washed with 0.1 % PBS-T (0.1 % v/v tween-20 in PBS; 3 × 5.0 mL, five min each at room temperature) followed by incubation with appropriate fluorophore or horse radish peroxidase (HRP) conjugated secondary antibodies for 1.0 h at room temperature. Membranes were washed with 0.1 % PBS-T (3 × 5.0 mL, five min each) and then either directly scanned using Amersham Typhoon fluorescence scanner or developed on photographic films after treatment with enhanced chemiluminescence reagents. At least two independent replicate experiments were performed for all the blots.

The primary and secondary antibodies employed, along with dilution and stock concentration, are given below:

Primary antibodies:
Anti-CD15s (CSLEX1; IgM, dilution 1:5000; stock concentration 0.5 mg/mL)
Anti-PSGL1 (clone KPL1; IgG, dilution 1:5000, stock concentration 0.5 mg/mL)
Anti-CLA (HECA452; IgM, dilution 1:1000, stock concentration 0.5 mg/mL)
Anti-β-actin (AC15, IgG, dilution 1:20000, stock concentration 2.2 mg/mL)

Secondary antibodies:

AlexaFluor488-conjugated anti-mouse IgM (1: 10000, stock concentration 2.0 mg/mL)
AlexaFluor488-conjugated anti-rat IgM (1:10000, stock concentration 0.5 mg/mL)
HRP-conjugated anti-mouse IgG (1:10000, stock concentration 0.8 mg/mL)
Cy5-conjugated donkey anti-mouse IgG (1:10000, stock concentration 1.4 mg/mL)
AlexaFluorplus488-conjugated anti-mouse IgG (1:10000, stock conc. 2.0 mg/mL)

*Effect of ManNAc analogues on expression of MAL-II and SNA epitopes*: HL-60 cells ($3.0 \times 10^5$ cells/mL in 10 mL complete medium in 10 cm diameter Petri dishes) were incubated with **D**, **1a**, **1b**, or **1d** (50 μM), along with untreated (**U**) controls. After 72 h, cells were harvested and lyzed in RIPA buffer containing PIC. Protein concentration in the soluble fraction was estimated using Bradford assay, subjected to 10 % SDS-PAGE, blotted onto nitrocellulose membranes (constant current 250 mA, 3 h, 4 °C). Membranes were blocked using 2 % w/v gelatin (5.0 mL) in 0.1 % PBS-T at room temperature. After 1h, membranes were incubated with either bMAL-II (1:2000, stock conc. 1.0 mg/mL) or bSNA (1:5000, stock conc. 2.0 mg/mL) in 0.1% PBS-T (5.0 mL) at room temperature. After 1 h, membranes were washed using 0.1 % PBS-T (6 × 5.0 mL, 5.0 min), followed by incubation with HRP-conjugated avidin (1:50000, stock conc. 2.0 mg/mL) in 0.1% PBS-T (5.0 mL). After 30 min, membranes were washed using 0.1 % PBS-T (6 × 5.0 mL, 5.0 min). Blots were developed using enhanced chemiluminescence substrate and photographic films. Blots using anti-β-actin (as mentioned above) were performed in parallel as loading controls. At least two independent replicate experiments were performed.

*Effect of ManNAc analogues on cell adhesion to E-selectin and L-selectin coated surfaces*: This assay was performed in 96-well polystyrene plates under sterile conditions. The plates were prepared freshly on the day of adhesion experiments. Wells were first coated with protein-G (200 ng in 50 µL PBS per well) for 1.0 h at room temperature (RT) followed by PBS washes (2 × 0.1 mL). Then either recombinant mouse E-selectin-Fc chimera or L-selectin-Fc chimera (20 ng in 50 µL PBS per well) was added to the wells and plate was left undisturbed for 1.0 h at RT. Wells were aspirated and washed with PBS (2 × 0.1 mL) followed by blocking with 1.0 % (w/v) BSA in 50 µL PBS per well for 1.0 h at RT. After blocking, the wells were washed with PBS (3 × 0.1 mL).

HL-60 cells ($3.0 \times 10^5$ cells per mL; 2.0 mL in six well plates) were incubated with 50 μM ManNAc analogues, harvested at 48 h, washed once with serum free RPMI medium (0.5 mL), and resuspended in serum free RPMI 1640 media containing 1.5 mM $CaCl_2$ ($4.0 \times 10^5$ cells/mL in 1.0 mL). The single cell suspension was added to each well ($2.0 \times 10^4$ cells in 50 µL media per well) and allowed to adhere for 1.0 h at 37 °C in a $CO_2$ incubator. Thereafter, the wells were washed with PBS (2 × 0.1 mL), cells were fixed with 4.0 % (w/v) paraformaldehyde (PFA) in PBS (50 μL) for 10 min followed by staining with DAPI (1:3000) in 50 µL PBS for 20 min after PBS washes (3 × 0.1 mL). Excess stain was removed with PBS (3 × 0.1 mL) and 50 μL PBS was added to each well. Wells coated and blocked with protein-G and BSA, respectively, (without selectins) served as negative controls. Images were captured at 10× magnification using Olympus IX53 fluorescence microscope. At least two fields were acquired from each well. For each condition, there were three replicate wells. The images were analyzed manually and the total number of adherent cells per well for each condition were plotted.

#### 3.2.4. Estimation of total sialic acids by periodate-resorcinol assay

Total sialic acids were measured following reported procedures.[5, 11] HL-60 cells ($3.0 \times 10^5$ cells/mL in 10 mL complete medium in 10 cm diameter Petri dishes) were incubated with

**D**, **1a**, **1b**, **1c**, **1d**, **1e**, **1f**, **1g**, **2a**, or **2b** (100 μM each), along with untreated (U) controls. After 48 h, cells were harvested, washed with PBS (3 × 5.0 mL; 3000×*g*, for 2.0 min), aliquoted to 2 × $10^6$ cells per sample and resuspended in PBS (150 μL) in 1.5 mL microcentrifuge tubes. Cells were lyzed by three freeze-thaw cycles and the total lysates were subjected to periodate-resorcinol assay. Briefly, lysates (150 μL) were treated with 0.4 M aqueous periodic acid (6.67 μM) or (2.5 μL) and incubated either on ice for 15 min (for measurement of total sialic acids) or at 37 °C for 1.0 h (for measurement of glycoconjugate bound sialic acids). About 10 mL of reagent cocktail was prepared using 6 % w/v of aqueous resorcinol (1.0 mL), 2.5 mM copper (II) sulfate (1.0 mL), 44 % of concentrated hydrochloric acid (4.4 mL), and water (3.6 mL). The mixture was then treated with the reagent cocktail (250 μL per sample), boiled at 100 °C for 10 min, rapidly quenched in an ice-bath and treated with *tert*-butanol (250 μL). The contents were centrifuged and the supernatants were aliquoted (200 μL) into 96-well plates and the absorbance at 630 nm was measured. At least three replicates were prepared for each condition.

#### 3.2.5. Immunoprecipitation of glycoproteins

HL-60 cell were incubated with ManNAc analogues **1a** – **1g** (50 μM, 72 h) and lyzed in RIPA buffer containing PIC (0.1 mL of lysis buffer was used for 10 × $10^6$ cells); Both untreated cells and cells treated with the vehicle (DMSO (D)) were included as controls in all experiments. The total cell lysates were subjected to immunoprecipitation using protein-G sepharose beads coated with (a) anti-PSGL-1 (clone KPL-1), (b) anti-CD43 (clone C-20, *C*-terminal) and (c) anti-CD44 (clone IM7) in order to isolate CD162/PSGL-1, CD43, and CD44 respectively.

*Preparation of beads*: Protein-G sepharose beads (30 μL of 50% slurry in 20% ethanol) in 1.5 mL tube were suspended in NP-40 buffer (0.1 % v/v Nonidet P-40 in PBS, pH 7.4, 1.0 mL), mixed well and centrifuged (200×*g*, 5.0 min at 4 °C), supernatant was aspirated, and the beads were re-suspended in NP-40 buffer (0.5 mL), treated with 1.0 μg of anti-PSGL-1 (clone KPL-1) antibody and allowed to bind for 3 h on an end-to-end rotor at 4 °C. Beads were centrifuged, supernatants were discarded, and the antibody-protein-G beads were washed with NP-40 buffer (2 × 0.5 mL).

*Immunoprecipitation of PSGL-1/CD162 and CD43*: Total cell lysates (2.0 mg) were added through the side of the tube and the total volume was made up to 0.5 mL using NP-40 buffer and incubated overnight at 4 °C on an end-to-end rotor. Samples were centrifuged, supernatant was discarded, and the antigen-antibody-protein-G beads were washed gently with NP-40 buffer (2 × 1.0 mL). The beads were then suspended in 5× Laemmli buffer (20 μL) and boiled at 100 °C for 8-10 min. Samples were cooled to room temperature, centrifuged, the supernatants were resolved on 5.0 % SDS-PAGE gels, blotted on to nitrocellulose membranes, blocked using non-fat milk in PBS (5.0 % v/v; 5.0 mL), and probed with either anti-PSGL-1 (dilution 1:2500) or anti-CD43 and anti-sLeX (clone CSLEX1) (dilution 1:5000) using independent blots. The blots were developed using appropriate HRP-conjugated secondary antibodies and signal were observed using enhanced chemiluminescence.

*Immunoprecipitation of CD44*: HL-60 cells were incubated with D, 1b, 1d, or 1g (50 mM, 72 h), then lysed in RIPA buffer (10 × $10^6$ cells in 100 μL) and protein concentrations were estimated using the Bradford assay. Total cell lysates (1.0 mg) were treated with anti-CD44 (clone IM7) antibody (1.0 μg) and gently agitated overnight at 4 °C. The immune complexes were then captured by the addition of protein-G agarose beads (30 μL of 50 % slurry; 20 mg/mL protein-G; pre-washed with PBS) and incubated at 4 °C for 2 h with gentle mixing. Samples were centrifuged (14000×*g*, 5.0 min), supernatants were discarded, and washed with

0.1% v/v NP-40 buffer (3 × 1.0 mL; 5.0 min of incubation for each wash). The beads were resuspended in 5x Laemmli sample buffer (30 μL), boiled for 10 min, resolved on SDS-PAGE, blotted, blocked with 5.0 % v/v non-fat milk in PBS, and probed by western blotting using biotinylated anti-CD44 (clone IM7) and anti-sLeX (clone CSLEX1) antibodies as given above.

#### 3.2.6. Statistical methods and softwares used in this study

For plotting graphs and calculation of significant probability ‘p’ values, one-way ANOVA, two-way ANOVA, student t-test, along with Bonferroni post-hoc correction, were employed in Microsoft Excel or GraphPad Prism 8.

### 4. List of reagents used in this study

| List of reagents and source | Source | Identifier |
|---|---|---|
| **Cell lines** | | |
| Jurkat | Dr. A. Qadri, NII | -- |
| HL-60 | ECACC | Cat. No. 98070106 |
| **Peracetyl HexNAc Analogues** | | |
| $Ac_4$ManNCp (**1a**) | In-house | |
| $Ac_4$ManNCb (**1b**) | -do- | |
| $Ac_4$ManNAz (**1c**) | -do- | |
| $Ac_4$ManNAc (**1d**) | -do- | |
| $Ac_4$ManNProp (**1e**) | -do- | |
| $Ac_4$ManNBut (**1f**) | -do- | |
| $Ac_4$ManNPent (**1g**) | -do- | |
| $Ac_4$GlcNCp (**2a**) | -do- | |
| $Ac_4$GlcNCb (**2b**) | -do- | |
| $Ac_4$GalNAz (**3**) | -do- | |
| **Antibodies** | | |
| Mouse anti-human CD15s (clone CSLEX1) | BD Biosciences | Cat. No. 551344 |
| Mouse IgM kappa isotype control | -do- | Cat. No. 555581 |
| Rat anti-human Cutaneous Lymphocyte Antigen (clone HECA-452) | -do- | Cat. No. 555946 |
| Rat IgM kappa isotype control | -do- | Cat. No. 555950 |
| Mouse anti-human CD15 (clone HI98) | -do- | Cat. No. 555400 |
| Alexa Fluor 488 anti-mouse IgM | Invitrogen | Cat. No. A21042 |
| Alexa Fluor 488 anti-rat IgM-Mu chain | -do- | Cat. No. A21212 |
| Goat anti-rat IgM Mu chain (HRP) | Abcam | Cat. No. ab98373 |
| R-Phycoerythrin AffiniPure $F(ab')_2$ fragment goat Anti-human IgG + IgM (H+L) | Jackson laboratories | Cat. No. 109-116-127 |
| Anti-β-actin antibody, mouse monoclonal | Sigma-Aldrich | Cat. No. A1978 |
| Peroxidase AffiniPure goat anti-mouse IgG (H+L) | Jackson laboratories | Cat. No. 115-035-003 |
| Cy5 AffiniPure donkey anti-mouse IgG (H+L) | -do- | Cat. No. 715-175-150 |

| | | |
|---|---|---|
| Goat anti-mouse IgG (H+L), AlexaFluor plus 488 | Invitrogen | Cat. No. A32723 |
| **Recombinant proteins and lectins** | | |
| Human E-Selectin-Fc chimera protein, Recombinant (His and hFc tag) | Sino biological | Cat. No. 10335-H03H |
| Recombinant mouse L-selectin-Fc (IgG1)-His$_6$ chimera protein | BioLegend | Cat. No. 772804 |
| Recombinant mouse E-Selectin (CD62E)-Fc(IgG1)-His$_6$-tag Chimeric (carrier-free) | BioLegend | Cat. No. 755504 |
| Biotinylated elderberry bark lectin (bSNA) | Vector laboratories | Cat. No. B-1305 |
| Biotinylated *Maackia amurensis* (bMAL II) | -do- | Cat. No. B-1265 |
| Avidin-Peroxidase Conjugate from hen egg white/horse radish | Sigma-Aldrich | Cat. No. 11371 |
| FITC-avidin | -do- | Cat. No. A2901 |
| **Chemicals and other resources** | | |
| Dimethyl sulfoxide (DMSO) | Emparta (Merck) | Cat. No. 1.07046.0521 |
| Propidium iodide (PI) | Sigma-Aldrich | Cat. No. P4170 |
| Dibenzocyclooctyne-Cy5 (DBCO-Cy5) | Click chemistry Tools | Cat. No. A130-25 |
| RPMI-1640 | Lonza | Cat. No. 12-702F |
| Fetal bovine serum (FBS) | Gibco | Cat. No. 10082-147 |
| Penicillin / streptomycin (P/S) | Gibco | Cat. No. 15070-063 |
| Skim milk powder | HiMedia | Ref. No. GRM1254 |
| Protein G | Pierce | Cat. No. 0021193 |
| Protease inhibitor cocktail (PIC) | Sigma Aldrich | Cat. No. P8340 |
| Sytox Red (5.0 mM in DMSO) | Life Technologies | Cat. No. S34859 |
| Protein-G sepharose | GE Healthcare | Cat. No. 17-0618-01 |
| Nonidet P-40 | Spectrochem | Cat. No. 011487 |

### 5. References

1. Hassenruck, J. & Wittmann, V. Cyclopropene derivatives of aminosugars for metabolic glycoengineering. *Beilstein J Org Chem* **15**, 584-601 (2019).
2. Saxon, E. & Bertozzi, C.R. Cell surface engineering by a modified Staudinger reaction. *Science* **287**, 2007-2010 (2000).
3. Shajahan, A. et al. Carbohydrate-Neuroactive Hybrid Strategy for Metabolic Glycan Engineering of the Central Nervous System in Vivo. *J Am Chem Soc* **139**, 693-700 (2017).
4. Jacobs, C.L. et al. Substrate specificity of the sialic acid biosynthetic pathway. *Biochemistry* **40**, 12864-12874 (2001).
5. Kim, E.J. et al. Characterization of the metabolic flux and apoptotic effects of O-hydroxyl- and N-acyl-modified N-acetylmannosamine analogs in Jurkat cells. *J Biol Chem* **279**, 18342-18352 (2004).
6. Hang, H.C., Yu, C., Kato, D.L. & Bertozzi, C.R. A metabolic labeling approach toward proteomic analysis of mucin-type O-linked glycosylation. *Proc Natl Acad Sci U S A* **100**, 14846-14851 (2003).
7. Sampathkumar, S.G., Li, A.V. & Yarema, K.J. Synthesis of non-natural ManNAc analogs for the expression of thiols on cell-surface sialic acids. *Nat Protoc* **1**, 2377-2385 (2006).
8. Medgyes, A., Farkas, E., Liptak, A. & Pozsgay, V. *Tetrahedron*, 4159-4178 (1997).
9. Kim, T.Y. & Davidson, E.A. Synthesis of acetylhexosamine-1-phosphate. *J. Org. Chem.* **28**, 2475-2476 (1963).
10. Agarwal, K. et al. Inhibition of mucin-type O-glycosylation through metabolic processing and incorporation of N-thioglycolyl-D-galactosamine peracetate (Ac5GalNTGc). *J Am Chem Soc* **135**, 14189-14197 (2013).
11. Jourdian, G.W., Dean, L. & Roseman, S. The sialic acids. XI. A periodate-resorcinol method for the quantitative estimation of free sialic acids and their glycosides. *J Biol Chem* **246**, 430-435 (1971).

### 6. NMR ($^1$H and $^{13}$C) spectra for Ac$_4$ManNCb (1b) and Ac$_4$GlcNCb (2b)

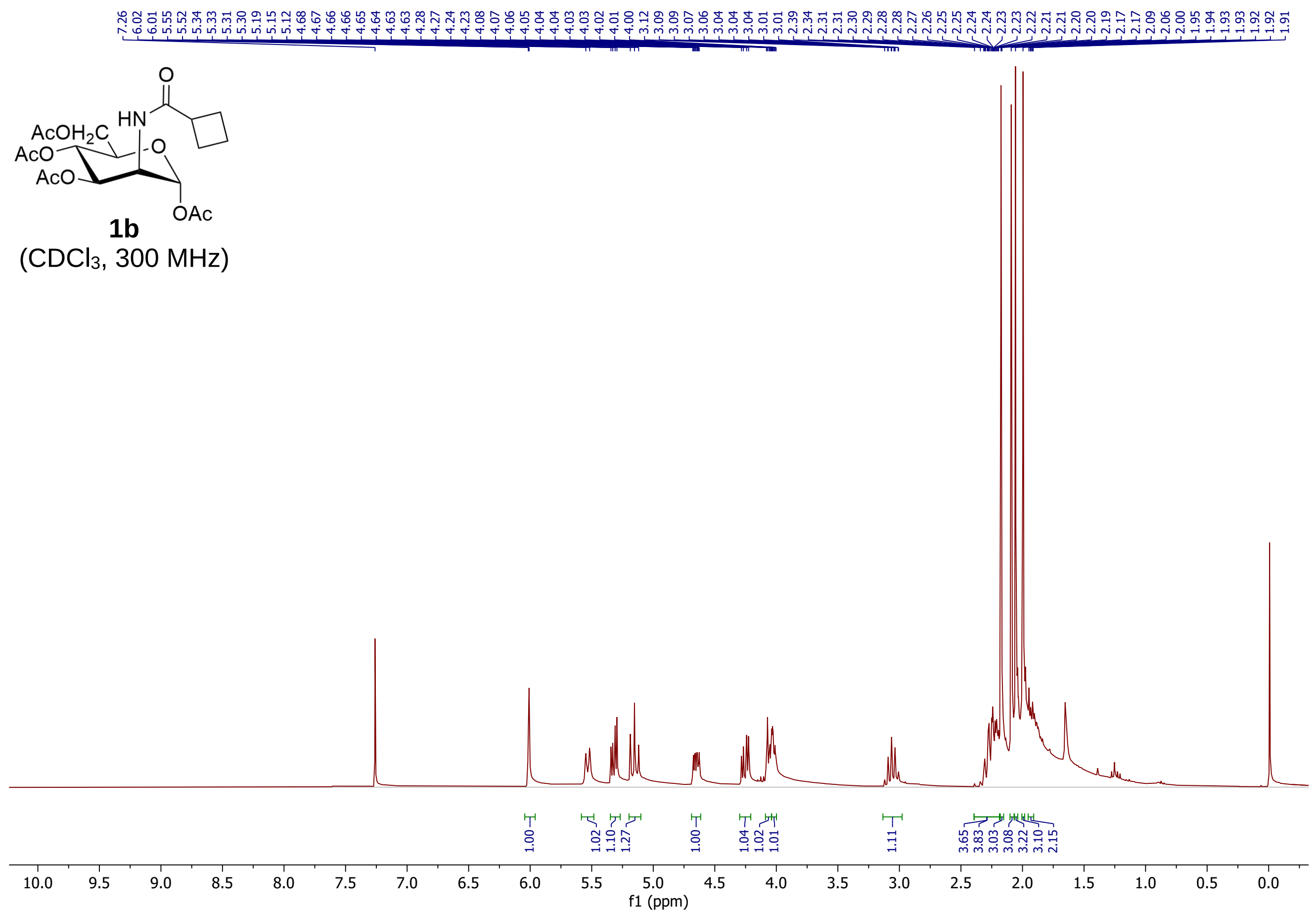

AcOH2C
AcO
AcO
HN
O
O
OAc
1b
(CDCl3, 300 MHz)
f1 (ppm)

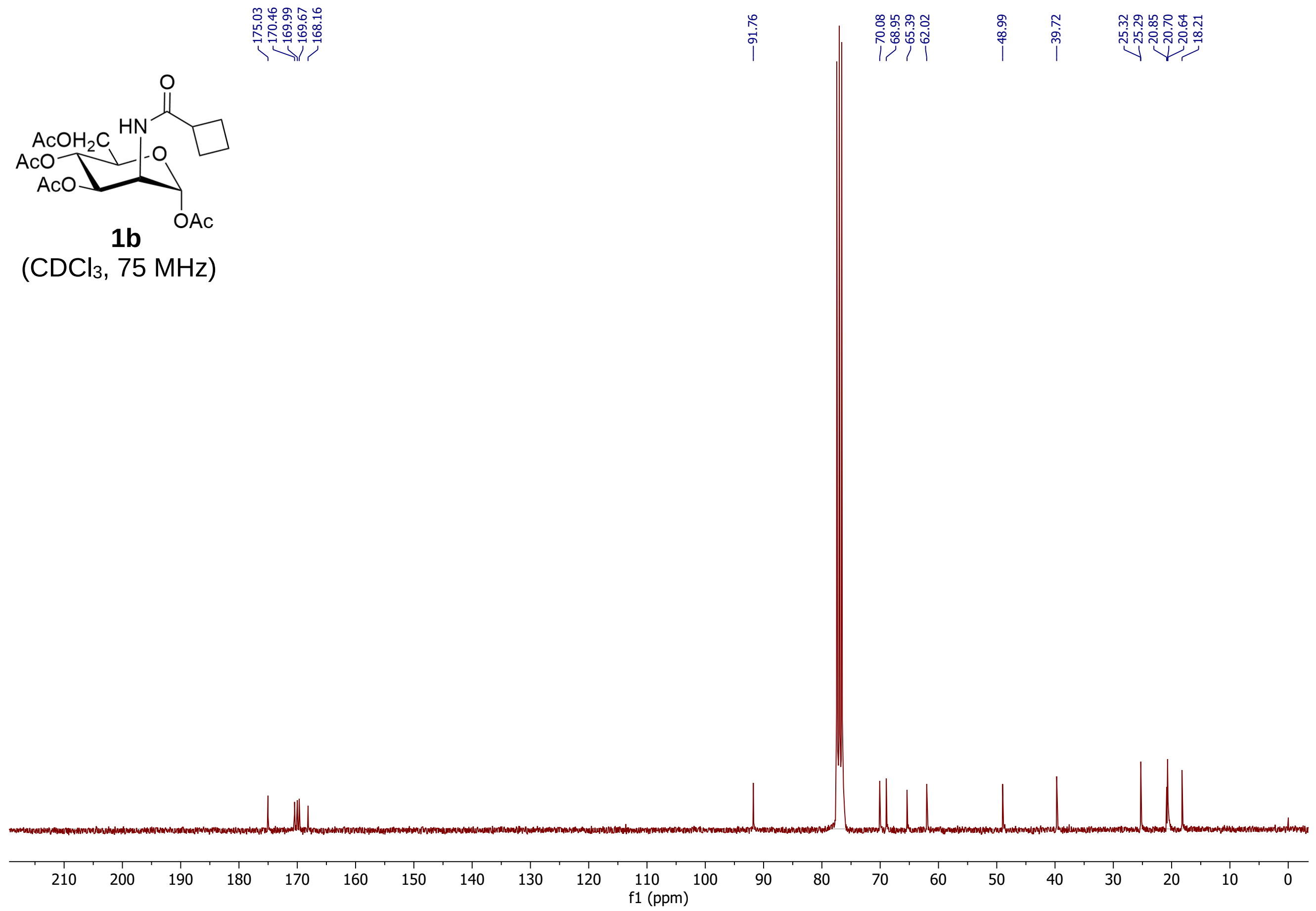

AcOH2C
AcO
AcO
HN
O
O
OAc
1b
(CDCl3, 75 MHz)
175.03
170.46
169.99
169.67
168.16
91.76
70.08
68.95
65.39
62.02
48.99
39.72
25.32
25.29
20.85
20.70
20.64
18.21
f1 (ppm)

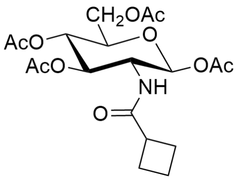

**2b**
(CDCl$_3$, 300 MHz)

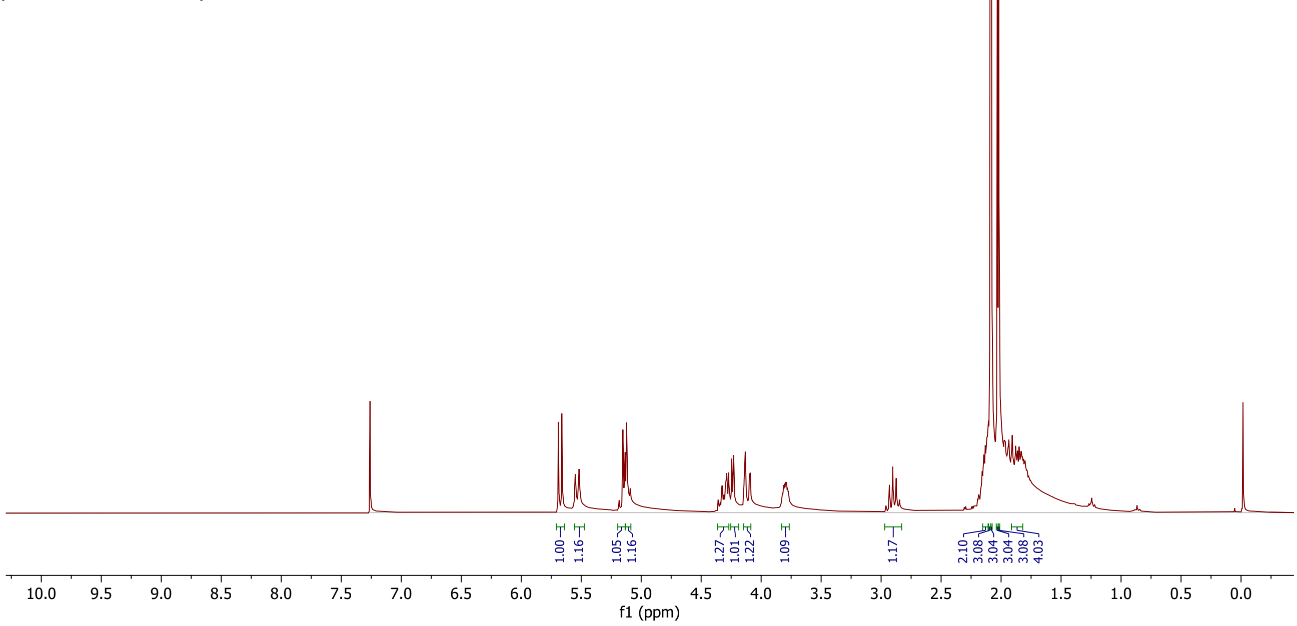

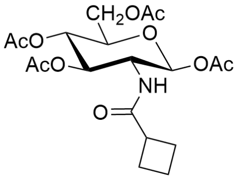

**2b**
(CDCl$_3$, 75 MHz)

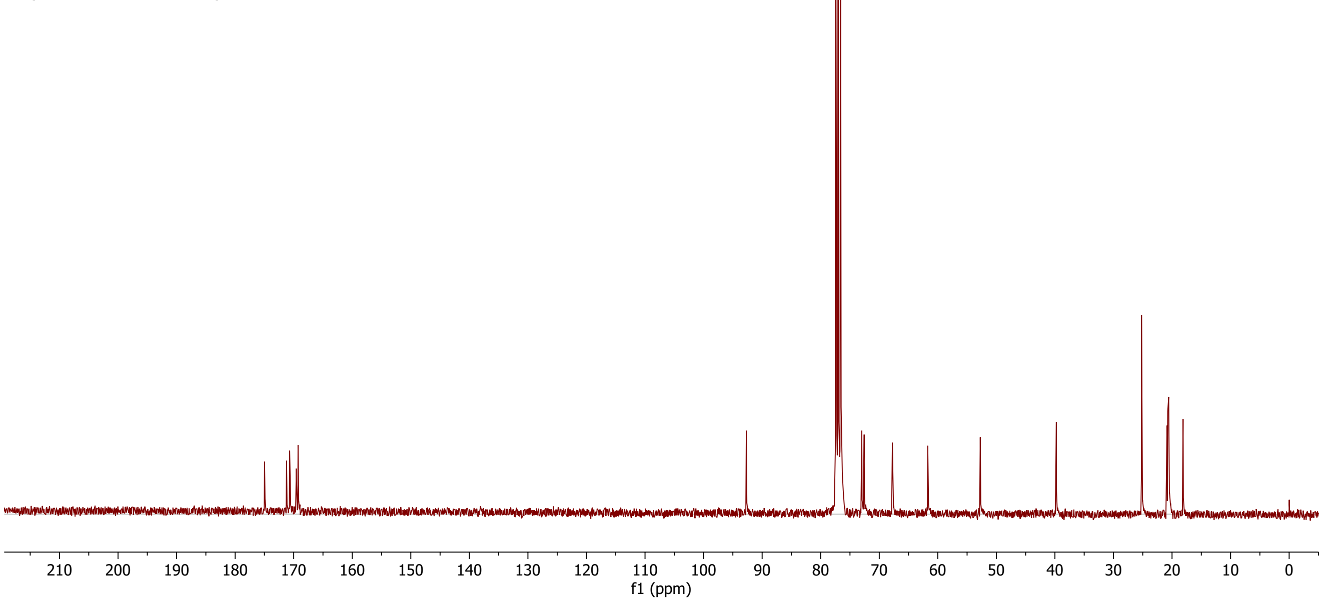

### 7. Supporting Figures 1 – 14

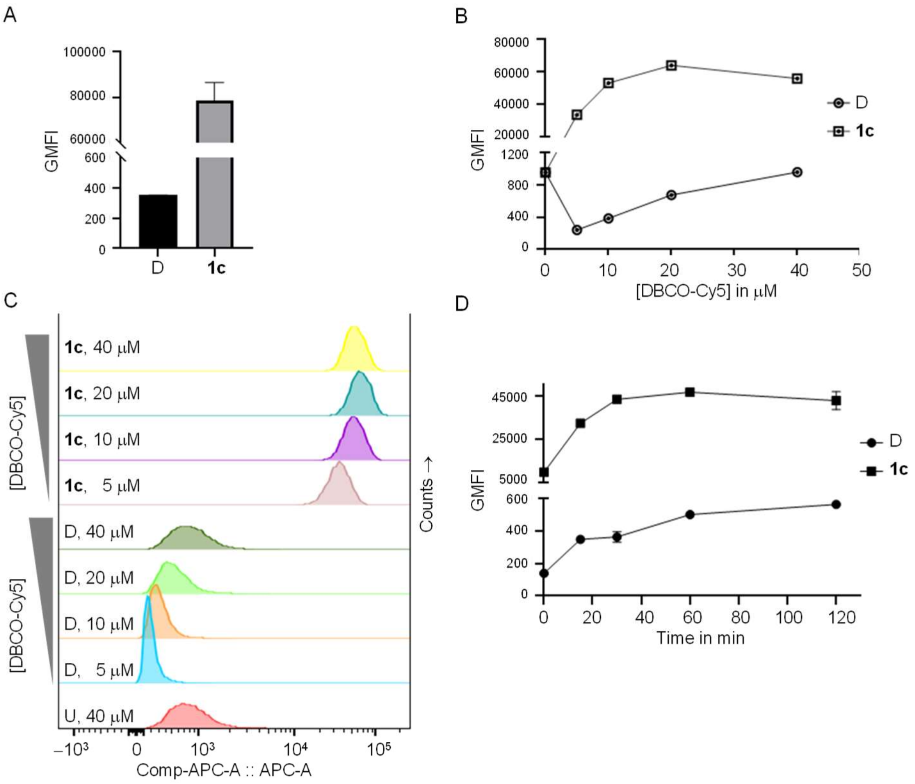

**Supporting Figure 1. Optimization of conditions for SPAAC in Jurkat cells**. (A) A bar graph showing the fluorescent intensity of Jurkat cells incubated with either D (vehicle) or **1c** (50 μM, 48 h) followed by click chemistry using DBCO-Cy5 (5.0 μM), 60 min). Dependency of SPAAC labeling on the concentration of DBCO-Cy5 (0 – 40 μM) incubated with Jurkat cells, pre-treated with either D or **1c** (50 μM, 24 h), for 60 min shown in (B) graphs and in (C) histograms. (D) Time course of SPAAC reaction of DBCO-Cy5 (20 μM) with Jurkat cells pre-treated with either D or **1c** (50 μM, 24 h). All samples were prepared in two replicates and 10,000 events of PI-negative populations were collected for each conditions by flow cytometry. SPAAC, strain-promoted azide-alkyne cycloaddition; GMFI, geometric mean fluorescent intensity.

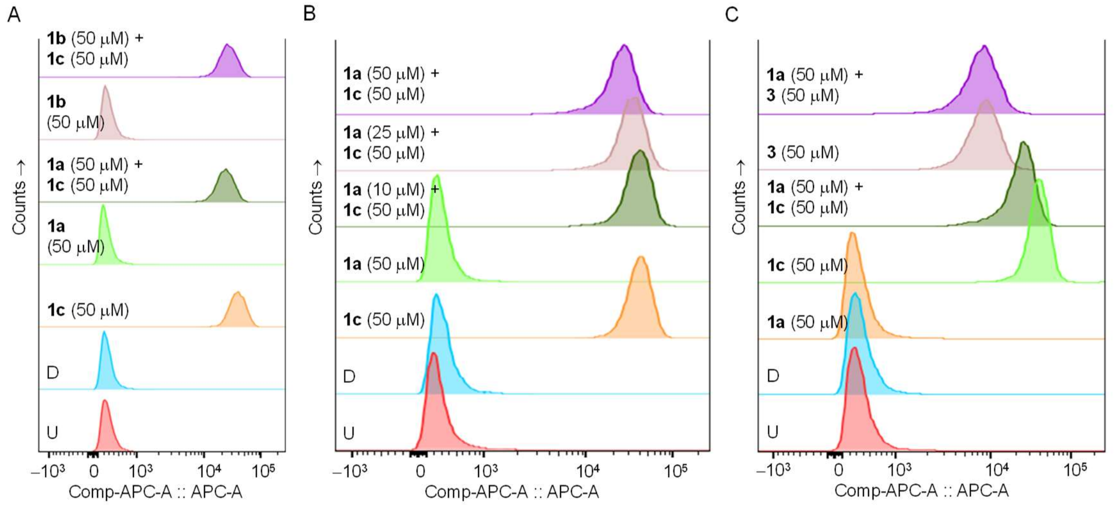

**Supporting Figure 2. Effect of HexNAc analogues treatment on NeuAz expression in Jurkat cells**. (A) Representative histograms showing fluorescent intensity levels in cells incubated, for 24 h, with either **1c** alone or in combination with **1a** or **1b** followed by DBCO-Cy5 reaction. Results show that both **1a** and **1b** competitively reduced the expression of NeuAz induced by **1c**. (B) Representative histograms showing fluorescent intensity levels in cells incubated with 1c alone or in combination with 1a at various concentrations, for 24 h, followed by DBCO-Cy5 reaction. Results showed that **1a**, reduced the NeuAz expression induced by **1c** in a dose-dependent manner. (C) Representative histograms showing fluorescent intensity levels in cells incubated alone with **1a**, **1c**, or **3** and in combination as **1a** with **1c** and **1a** with **3**. Results show that the NeuAz expression induced by **1c**, but not the GalNAz expression induced by **3**, is reduced by treatment with **1a**.

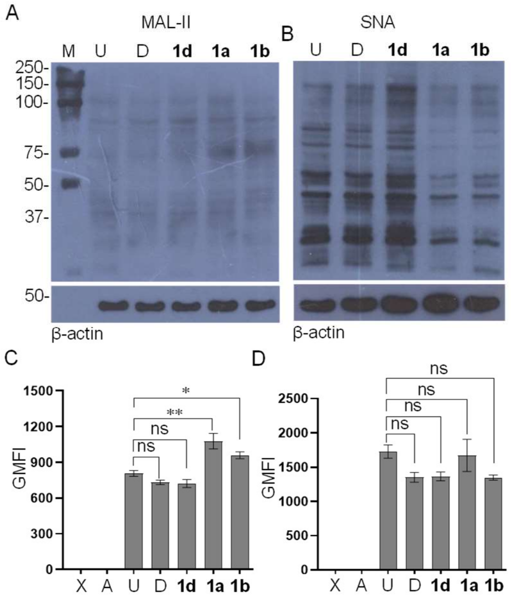

**Supporting Figure 3. Profiling of global effects on glycosylation using lectins**. HL-60 cells were incubated with D, **1a**, **1b**, or **1d** (50 μM, 72 h), lysed, resolved by SDS-PAGE, blotted on to nitrocellulose membrane and probed with (A) biotinylated *Maackia amurensis* lectin-II (bMAL-II) and (b) biotinylated *Sambucus nigra* agglutinin (bSNA) followed by horse radish peroxidase (HRP)-conjugated avidin. B-actin blots were employed as loading controls. Blots shown are representative of at least two biological replicates. HL-60 cells incubated with D, **1a**, **1b**, or **1d** (50 μM, 24 h) were harvested, stained with (C) bMAL-II and (D) bSNA followed by FITC-conjugated avidin and analyzed by flow cytometry. Errors bars shown are standard deviation of at least two biological replicate samples, each counted twice. M, molecular weight markers; U, untreated; D, DMSO (vehicle); X, unstained cells; A, FITC-avidin alone; GMFI, geometric mean fluorescence intensity.

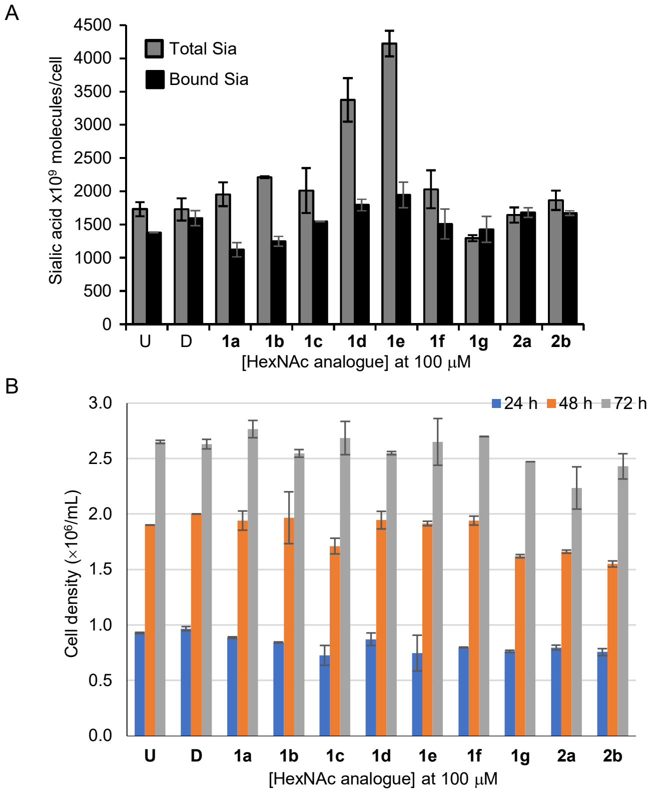

**Supporting Figure 4. Effect of HexNAc analogues on sialic acid content and cell density**. (A) Bar graph showing the levels of total sialic acids and glycoside bound sialic acids in HL-60 cells incubated with the HexNAc analogues (100 μM, 48 h) as measured by the periodate-resorcinol assay. Error bars shown as standard deviation of two replicate samples. Two independent experiments were performed. (B) Bar graph showing the densities at 24, 48, and 72 h of HL-60 cells incubated with HexNAc analogues (100 μM). Error bars shown are standard deviation of two replicate cell counts as measured by the Z2 particle counter.

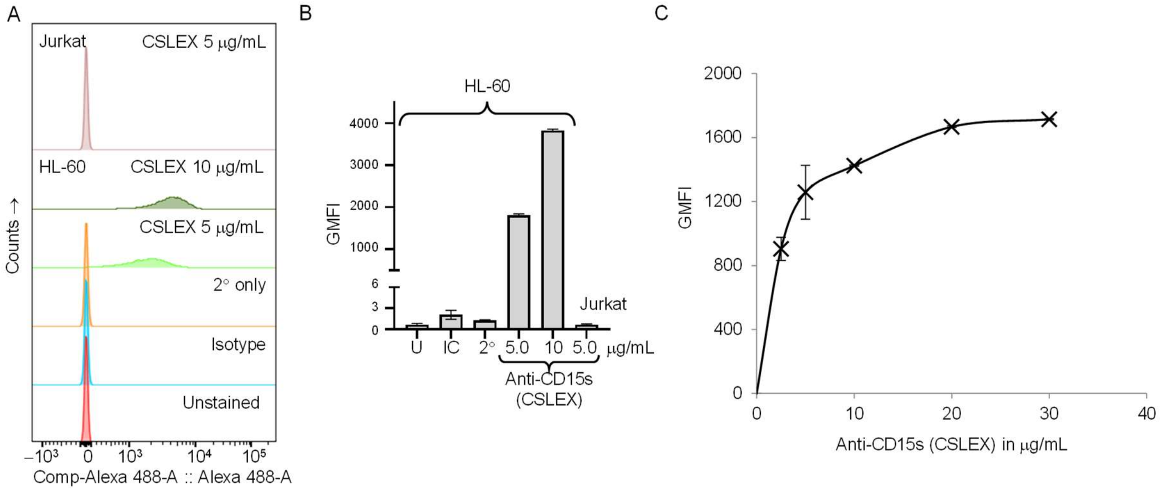

**Supporting Figure 5. Estimation of sialyl-Lewis X (CD15s) epitopes**. (A) Representative histograms showing fluorescent intensities of HL-60 cells stained with anti-CD15s (CSLEX1) antibody followed by AlexaFluor488-conjugated secondary antibody. Unstained, isotype antibody stained, and secondary-antibody alone controls are shown. Jurkat cells were found to be negative for CD15s (upper histogram). (B) Bar graphs showing the fluorescent intensity in HL-60 and Jurkat cells, along with controls. (C) Titration of saturation of CSLEX1 binding to HL-60 cells incubated with various concentrations of the CSLEX1 antibody. The error bars shown are standard deviation of duplicate samples. About 10000 events of PI-negative populations were counted by flow cytometry. GMFI, geometric mean fluorescence intensity.

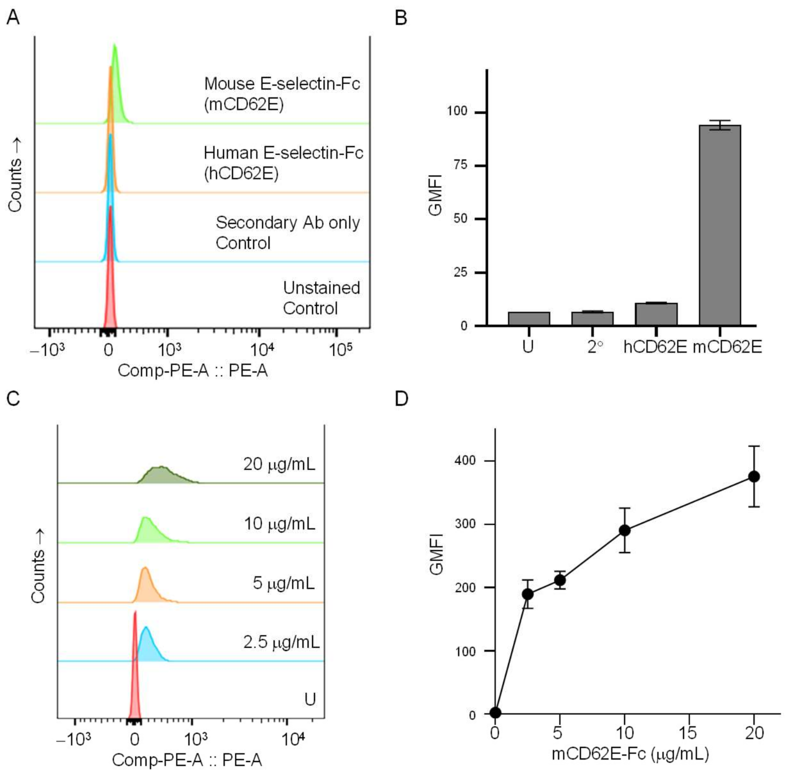

**Supporting Figure 6. Binding of E-selectin-Fc to HL-60 cells**. (A) Histograms and (B) bar graphs showing the fluorescence intensities of HL-60 cells stained either with human-E-selectin-Fc (hCD62E-Fc) or mouse-E-selectin-Fc (mCD62E-Fc) chimera proteins followed by secondary anti-Fc antibody. (C) Histograms and (D) graphs showing the fluorescent intensities of HL-60 cells stained with various concentrations of mCD62-Fc chimera. Samples were prepared in duplicates and 10000 events of Sytox Red -negative population were counted. Error bars shown are standard deviations.

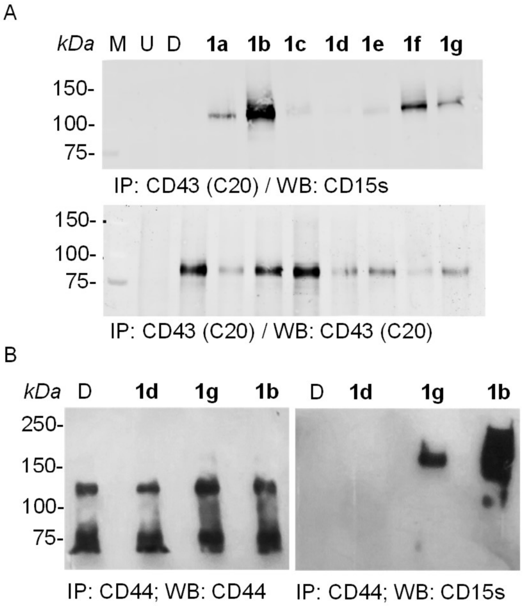

**Supporting Figure 7. ManNAc analogues 1b enhances CSLEX1 epitopes on CD43 and CD44**. HL-60 cells were incubated with the ManNAc analogues 1a – 1g (50 mM, 72 h) and lyzed. The total lysates were subjected to immunoprecipitation using protein-G agarose beads coated with (A) anti-CD43 (clone C-20, *C*-terminal) and (b) anti-CD44 (clone IM7). The immunoprecipitates were probed for the presence of sLeX epitopes using the anti-sLeX (clone CSLEX1) antibody by western blotting. Western blots for CD43 and CD44 are shown as controls for immunoprecipitation. Blots shown are representative of at least two biological replicates.

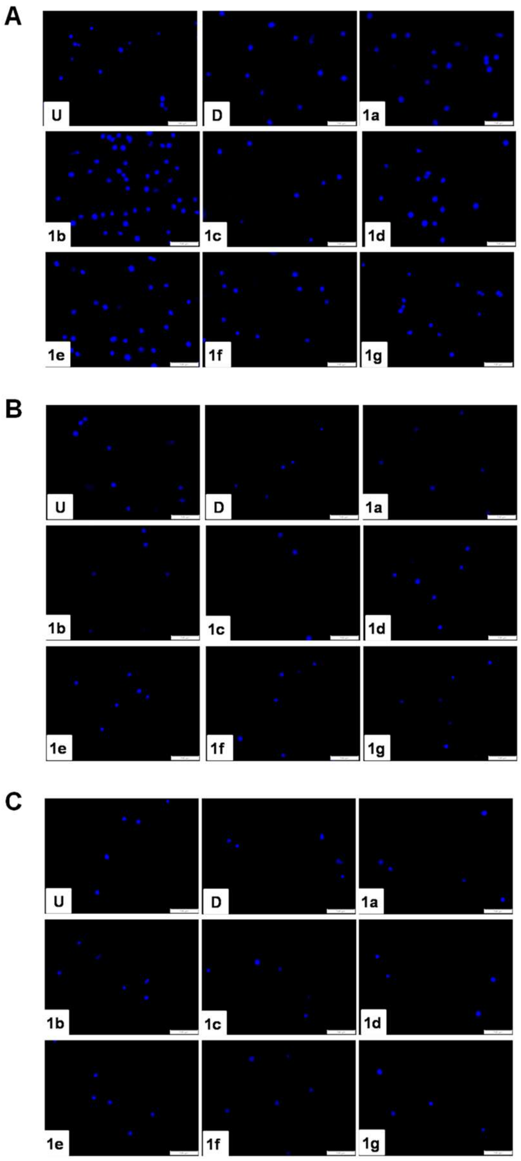

**Supporting Figure 8. Processing of ManNAc analogues modulates adhesion to selectins**. HL-60 cells were treated with 1a – 1g (50 μM, 48 h), along with untreated (U), and vehicle (D) treated controls, and allowed to adhere on plastic surfaces pre-coated with A) mouse E-selectin-Fc/protein-G/BSA , B) mouse L-selectin-Fc/protein-G/BSA, and C) protein-G/BSA (control). Cells were fixed, stained with DAPI, and imaged using a fluorescence microscope. Images shown are representative of at least three replicates. Scale bar, 100 μm.

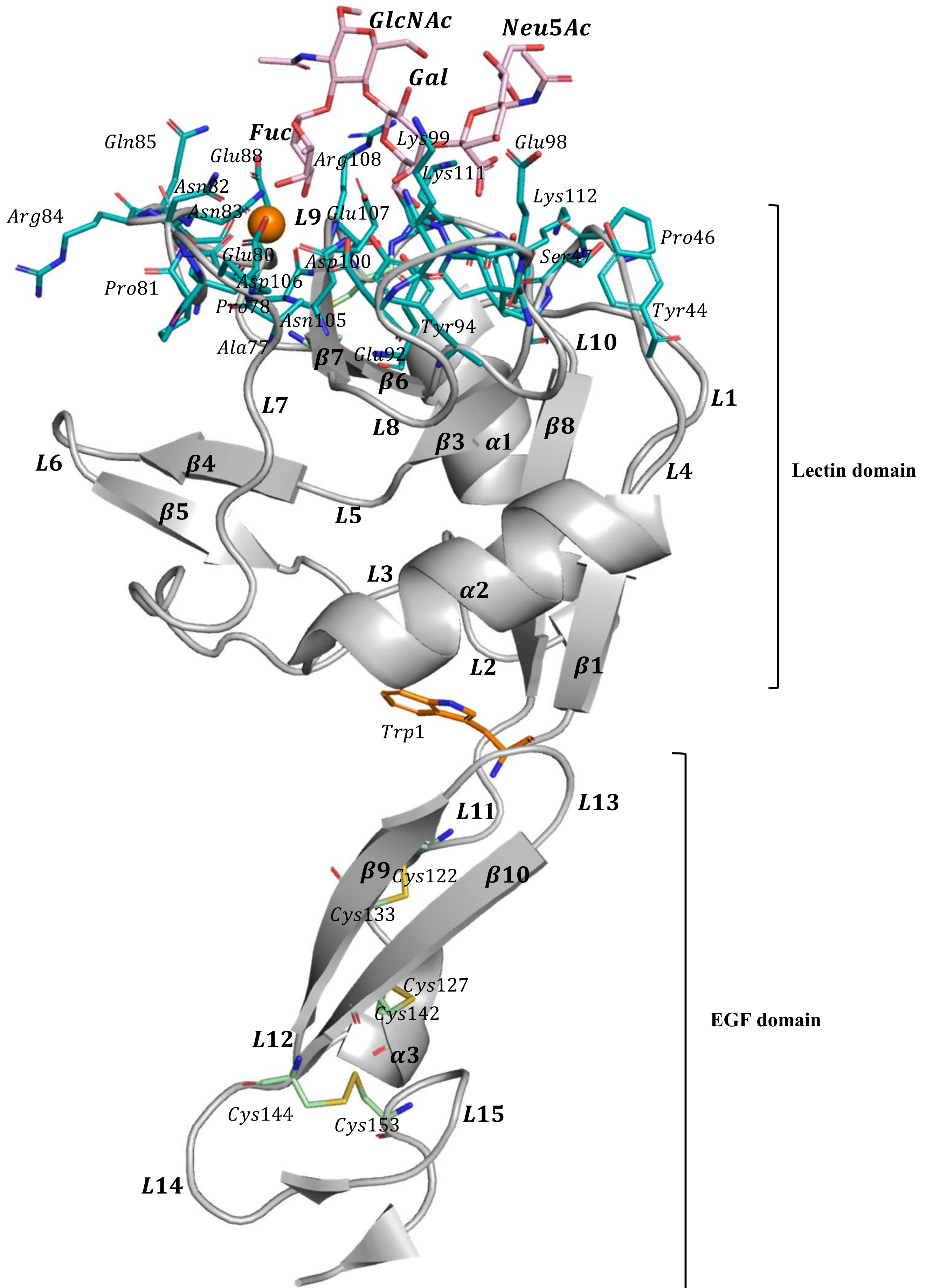

**Supporting Figure 9.** Structure of E-Selectin sLeX complex taken from PDB 1G1T. Shown in cartoon is the structure of E-Selectin (represented in grey cartoon) in complex with **sLeX** (shown in light pink sticks). The Ca2+ ion is shown in orange spheres. All the 10 cysteine residues that are part of 5 disulfide bonds (two in lectin domain and three in EGF domain) are shown in green and yellow sticks. Residues of ligand binding pocket are shown in teal sticks. The pivot residue (Trp1) is shown in orange sticks.

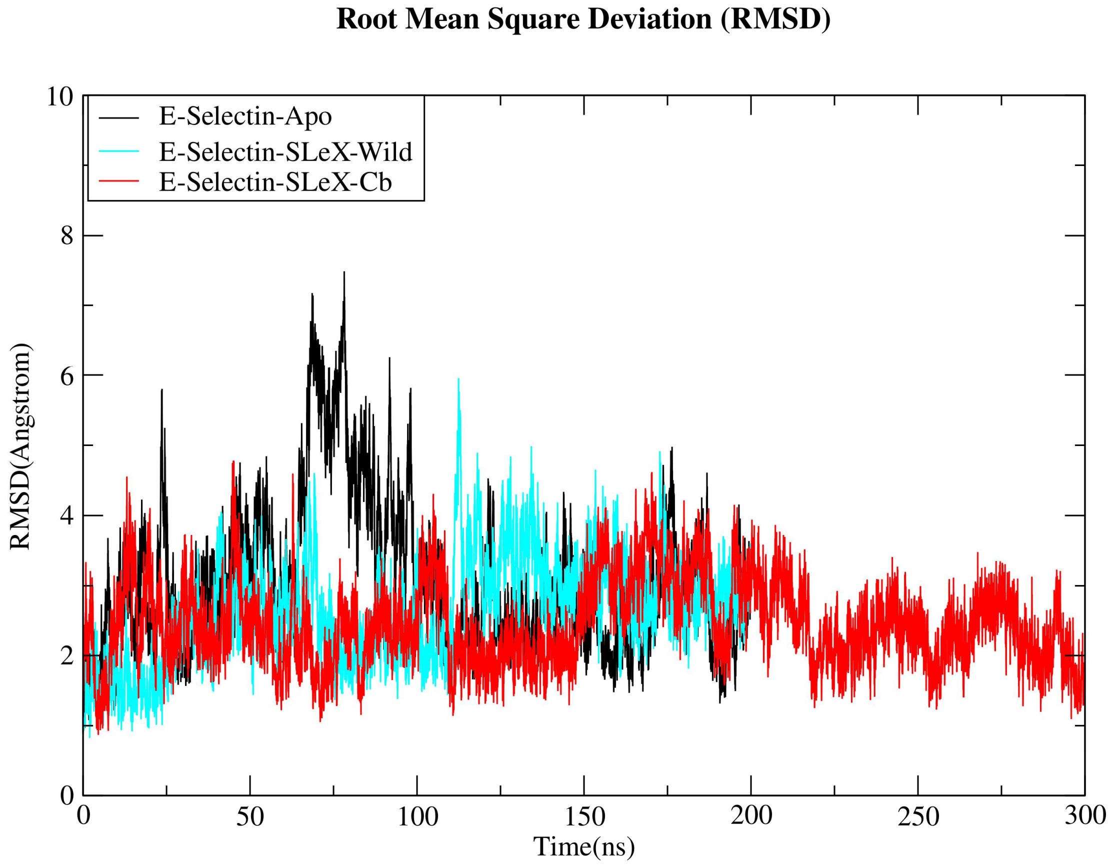

**Supporting Figure 10. Root Mean Square Deviation (RMSD).** Shows the RMSD of Cα atoms of E-Selectin, taken from all the three MD simulations with respect to time. RMSD of E-Selectin from **sLeX, sLeX-Cb complex** and apo (without the ligand) are shown in cyan, red and black respectively.

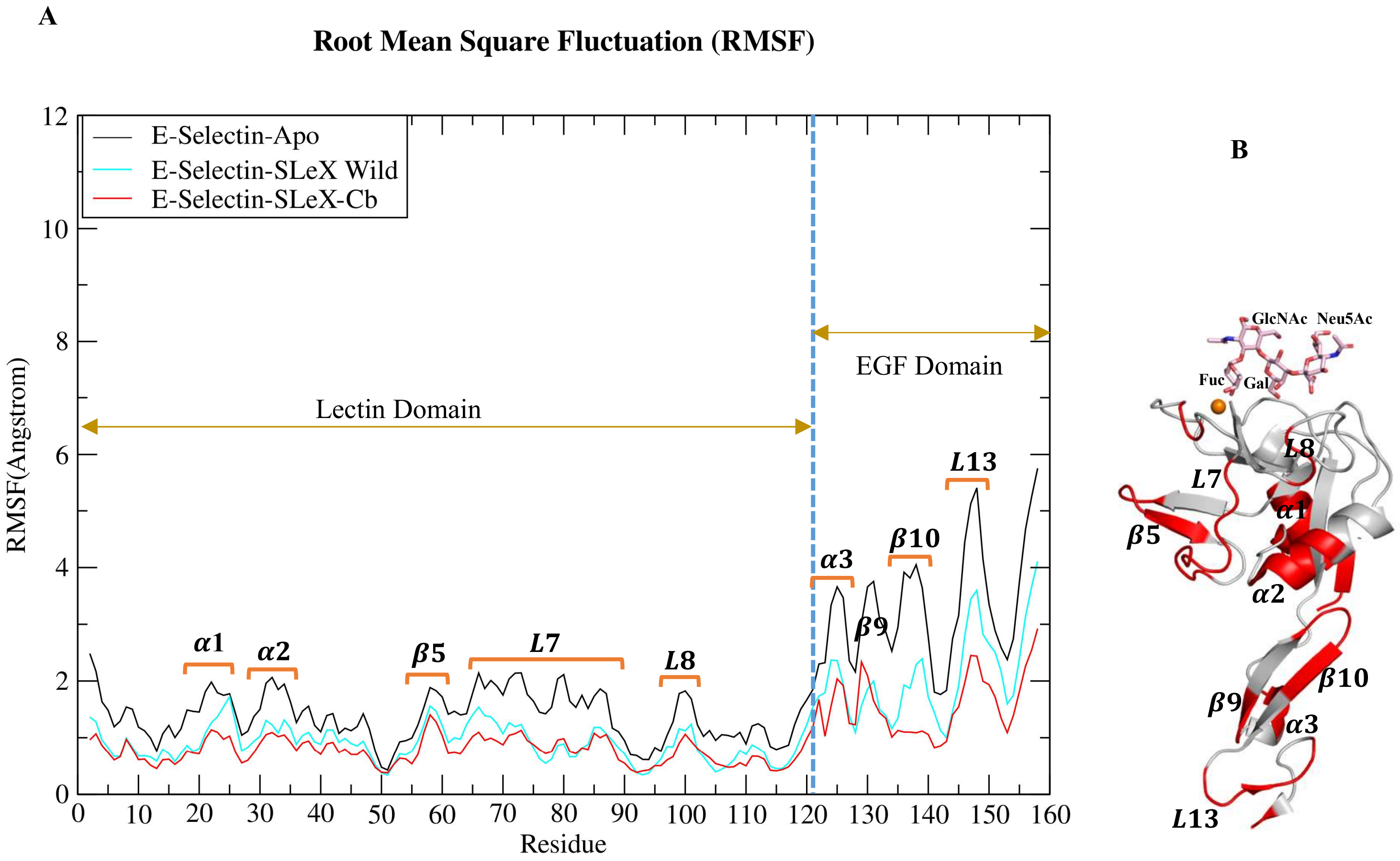

**Supporting Figure 11.** Shows Root Mean Square Fluctuations calculated for Cα atoms of E-Selectin taken from apo form as well as in complex with **sLeX** and its analog (**sLeX-Cb**). B) Shows the mapping of higher RMSF onto the structure of E-Selectin.

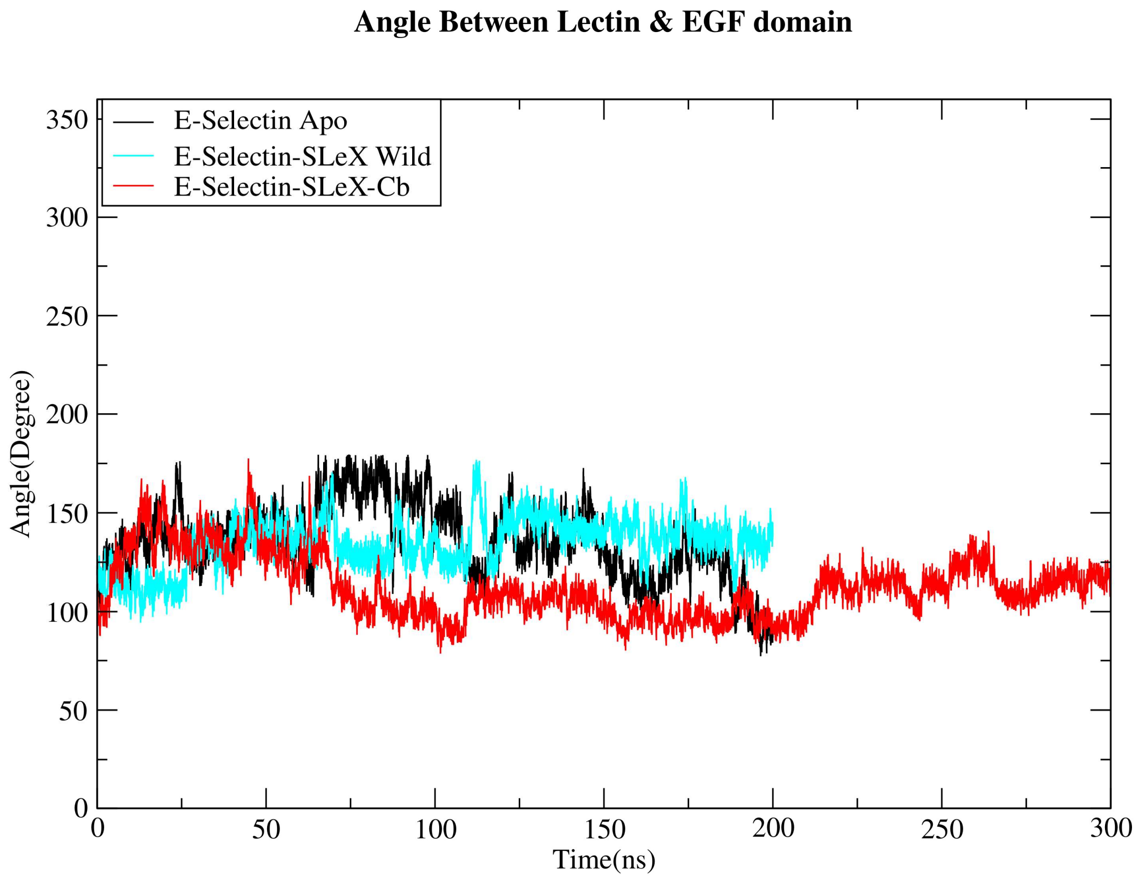

**Supporting Figure 12. Angle between the Lectin and EGF domain.** Plot showing the time evolution of interdomain angle between the lectin and EGF domain. Angle was calculated by taking the vectors connecting Calcium ion, C-α atom of Trp1 and Cys144 of E-Selectin. Black represents the interdomain angle of E-selectin taken from apo simulations, Cyan and red represents the interdomain angle of E-selectin in the presence of ligands **sLeX** and **sLeX-Cb** taken from complex simulations.

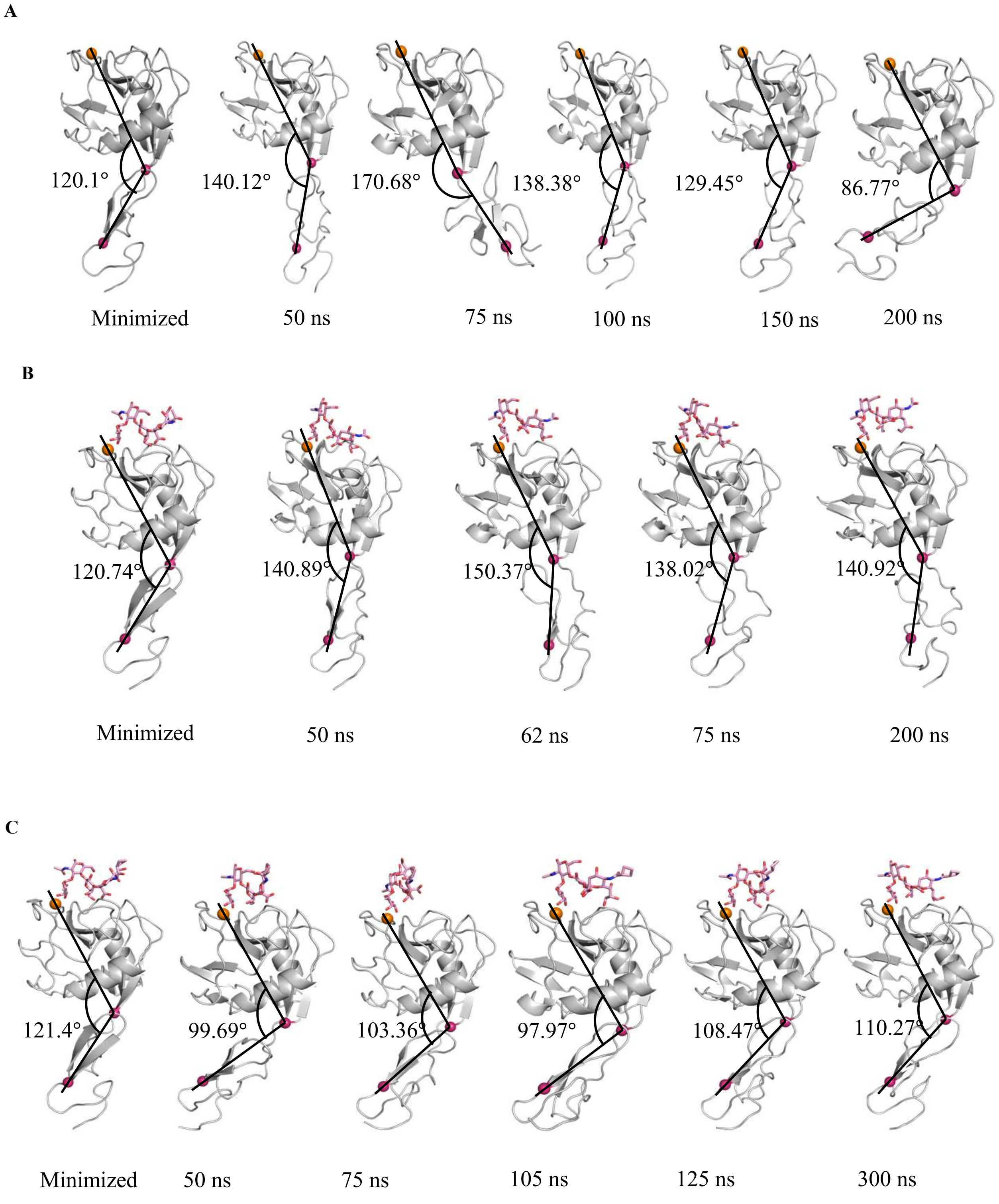

**Supporting Figure 13. Snapshots depicting the interdomain angle between the Lectin and EGF domain. A-C**) Snapshots showing the variation in the interdomain angle taken from the E-Selectin of apo (A) and **sLeX (B)** wild and **sLeX-Cb (C)** complex MD simulations. Angle was calculated by taking angle between the vectors connecting Calcium ion (orange sphere), C-α atoms from Trp1 and Cys144 (Winebrick spheres).

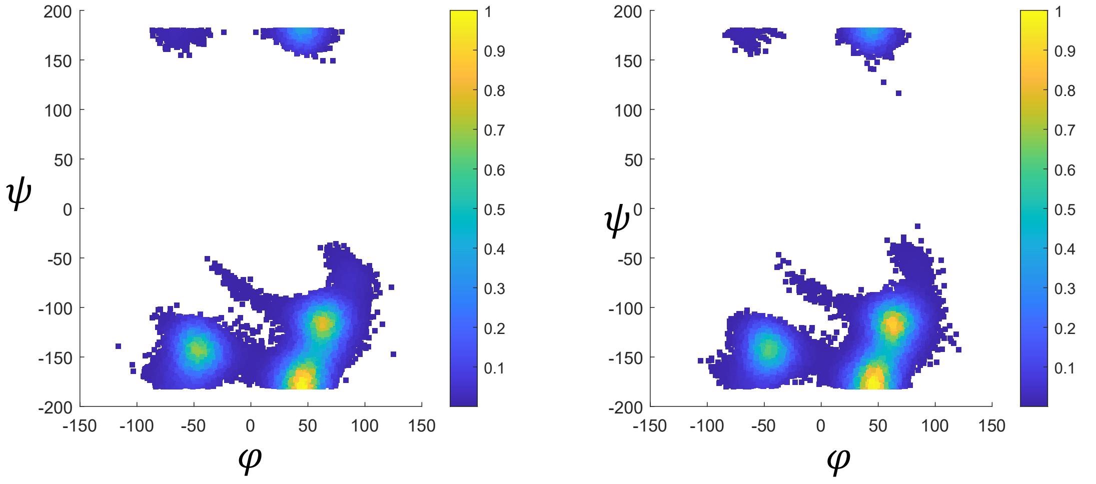

A B

**Supporting Figure 14. Ramachandran plot for dihedral angles between Neu5Acα2→3Gal of sLeX and sLeX-Cb.** Plots showing the distribution for (A) sLeX and (B) sLeX-Cb is the distribution for **sLeX-Cb** taken from the independent ligand MD simulations.

**Supporting Table 1. The mean RMSD and Standard deviation of C-$\alpha$ atoms for E-Selectin taken from the complex simulations of sLeX and its analog sLeX-Cb. Also, the mean RMSD and standard deviation of all atoms of ligands, calculated for all the snapshots taken from the complex and independent simulations are given.**

| S.No. | MD System | Average RMSD (Å) |
|---|---|---|
| 1. | E-Selectin Apo (200ns) | 3.09 $\pm$ 1.06 |
| 2. | E-Selectin- sLeX complex (200ns) | 2.62 $\pm$ 0.73 |
| 3. | E-Selectin- sLeX-Cb complex (300ns) | 2.45 $\pm$ 0.61 |
| 4. | sLeX alone (taken from the complex simulations) | 2.57 $\pm$ 0.34 |
| 5. | sLeX-Cb alone (taken from the complex simulations) | 2.11 $\pm$ 0.54 |
| 6. | sLeX Wild Alone ( 200ns ) | 1.87 $\pm$ 0.52 |
| 7. | sLeX-Cb Alone (200 ns) | 2.2 $\pm$ 0.59 |

**Supporting Table 2. Average dihedral angles calculated for all the three glycosidic linkages for sLeX wild and sLeX-Cb for the snapshots taken from the trajectory of independent ligand MD simulations.**

| S.No. | Glycosidic Linkage | Phi ($\phi$) | | Psi ($\psi$) | |
|---|---|---|---|---|---|
| | | sLeX – Wild | sLeX – Cb | sLeX – Wild | sLeX – Cb |
| 1. | **Fuc1→3GlcNAc** | -69.27° ± 8.9 | -69.16° ± 8.91 | -95.87° ± 7.54 | -95.88° ± 7.54 |
| 2. | **Gal1→4GlcNAc** | -68.46 ° ± 8.15 | -68.4° ± 8.12 | 130.4° ± 6.89 | 130.2° ± 6.85 |
| 3. | **Neu5Ac2→3Gal** | **31.4 ° ± 45.75** | **35.27° ± 42.23** | -122.5° ± 74.83 | -120.7° ± 81.23 |

**Supporting Table 3. The mean and standard deviation of all the dihedral angles of three glycosidic linkages of sLeX and sLeX-Cb of E-Selectin-sLeX/sLeX-Cb complexes, calculated for the snapshots taken whole MD simulation trajectory.**

| S.No. | Glycosidic Linkage | Phi ($\phi$) | | Psi ($\psi$) | |
|---|---|---|---|---|---|
| | | sLeX – Wild | sLeX – Cb | sLeX – Wild | sLeX – Cb |
| 1. | **Fuc1→3GlcNAc** | -72.47° ± 8.22 | -72.65° ± 8.53 | -95.31° ± 7.24 | -94.44° ± 7.47 |
| 2. | **Gal1→4GlcNAc** | -70.88 ° ± 8.18 | -73.91° ± 9.23 | 130.2° ± 6.49 | 132° ± 7.23 |
| 3. | **Neu5Ac2→3Gal** | **-37.2 ° ± 21.35** | **23.28° ± 50.77** | -139.6° ± 19.22 | -125.5° ± 32.32 |

**Supporting Table 4. Binding energetics (Kcal/mol) of E-Selectin- sLeX and E-selectin- sLeX-Cb complexes calculated for the snapshots taken from last 100 ns (100 – 200 ns in sLeX and 200 – 300 ns in sLeX-Cb complex) of MD trajectories, taken in four clusters.**

| S.No. | Complex | Energy Components | Cluster-1(100-125 ns) Complex | Cluster-2(125-150ns) Complex | Cluster-3(150-175ns) Complex | Cluster-4(175-200ns) Complex |
|---|---|---|---|---|---|---|
| **1.** | **E-Selectin – sLeX** | ΔvdW | -1233.77 | -1232.74 | -1234.07 | -1234.73 |
| | | $\Delta E_{EL}$ | **-12823.22** | **-12858.89** | **-12914.84** | **-12883.67** |
| | | $\Delta E_{GB}$ | -1991.44 | -1959.93 | -1915.83 | -1965.36 |
| | | $\Delta E_{SURF}$ | 63.16 | 62.52 | 62.39 | 62.68 |
| | | $\Delta G_{gas}$ (ΔvdW + $\Delta E_{EL}$) | -14056.99 | -14091.63 | -14148.92 | -14118.39 |
| | | $\Delta G_{solv}$ ($\Delta E_{GB}$+ $\Delta E_{SURF}$) | -1928.27 | -1897.41 | -1853.44 | -1902.67 |
| | | TOTAL (ΔH) ($G_{gas}$ + $G_{solv}$) | -15985.26 | -15989.04 | -16002.35 | -16021.06 |
| | | $T\Delta S_{Trans}$ | 16.49 | 16.49 | 16.49 | 16.49 |
| | | $T\Delta S_{Rot}$ | 16.86 | 16.87 | 16.86 | 16.87 |
| | | $T\Delta S_{Vib}$ | 77.24 | 74.50 | 75.75 | 74.13 |
| | | TOTAL (TΔS) | 110.59 | 107.87 | 109.11 | 107.49 |
| | | **ΔG (ΔH - TΔS)** | **-16095.85** | **-16097.01** | **-16111.47** | **-16128.55** |
| **2.** | **E-Selectin – sLeX-Cb** | | **Cluster-1(200-225 ns) Complex** | **Cluster-2(225-250ns) Complex** | **Cluster-3(250-275ns) Complex** | **Cluster-4(275-300ns) Complex** |
| | | ΔvdW | -1244.08 | -1247.84 | -1247.53 | -1248.25 |
| | | $\Delta E_{EL}$ | **-12971.12** | **-12951.82** | **-12951.55** | **-12955.10** |
| | | $\Delta E_{GB}$ | -1935.48 | -1942.41 | -1966.49 | -1946.62 |
| | | $\Delta E_{SURF}$ | 62.24 | 61.89 | 62.28 | 62.27 |
| | | $\Delta G_{gas}$ (ΔvdW + $\Delta E_{EL}$) | -14215.19 | -14199.66 | -14199.08 | -14203.35 |
| | | $\Delta G_{solv}$ ($\Delta E_{GB}$+ $\Delta E_{SURF}$) | -1873.24 | -1880.51 | -1904.21 | -1884.35 |
| | | TOTAL (ΔH) ($G_{gas}$ + $G_{solv}$) | -16088.43 | -16080.17 | -16103.29 | -16087.7 |
| | | $T\Delta S_{Trans}$ | 16.49 | 16.49 | 16.49 | 16.49 |
| | | $T\Delta S_{Rot}$ | 16.82 | 16.83 | 16.84 | 16.84 |
| | | $T\Delta S_{Vib}$ | 75.62 | 75.24 | 75.33 | 74.94 |
| | | TOTAL (TΔS) | 108.94 | 108.57 | 108.66 | 108.27 |
| | | **ΔG (ΔH - TΔS)** | **-16197.37** | **-16188.74** | **-16211.95** | **-16195.97** |
| | | Difference in ΔG (sLeX -Wild – sLeX-Analog) | -101.52 | -91.73 | -100.48 | -67.42 |
